# Sex differences in sRNA mediated transposon regulation under thermal stress in zebrafish germ cells

**DOI:** 10.1101/2025.11.15.688619

**Authors:** Alice M. Godden, Jean-Charles De Coriolis, Ruth Sullivan, Chelsea Drake, Simone Immler

## Abstract

Climate change-induced thermal stress threatens germline genome stability and triggers a molecular response of small RNA pathways involved transposable element (TE) regulation. The difference in mitotic and meiotic activity in male and female germ cells suggests that this response differs, but the mechanisms are elusive. Here we integrate whole-genome and transcriptome sequencing, and small RNA profiling to examine how mild thermal elevation affects TE regulation in male and female zebrafish gonads. We find that ovaries maintain genomic integrity through coordinated upregulation of piRNAs and miRNAs linked to heat-shock responses, limiting TE mobilisation. In contrast, testes exhibit reduced small RNA abundance, widespread TE activation, and increased *de novo* insertions, reflecting compromised genome defence. Multi-omic network analyses reveal a piRNA-TE-miRNA regulatory network preserved in ovaries but disrupted in testes under thermal stress. These findings reveal a sex-specific divergence in genome defence mechanisms, with implications for fertility and heritable genome stability in changing environments.

## Introduction

Climate change is increasing thermal variability through both short-term fluctuations and sustained warming, with important consequences for biological systems. Reproduction is particularly sensitive to temperature, and even transient thermal stress can impair fertility^1^. Environmental thermal variation triggers a molecular response in the germ cells which perturbs small RNA (sRNA) pathways and disrupts transposable element (TE) silencing, leading to TE de-repression, altered regulatory states, and increased transposition^2–5^. These responses can be transient epigenetic reprogramming, or persistent if TE insertions are retained in the genome. Central to this process are Piwi-interacting RNAs (piRNAs), which act as a primary defence against transposons and play critical roles in maintaining germline genome stability and regulating gene expression across generations^2–4,6^. While stress-induced TE activation is well documented in a range of taxa including plants^7^, insects^8^ and other invertebrates^9^ and vertebrates^5^, it remains unclear how these molecular responses in the germ cells are coordinated across regulatory levels, and whether TE mobilisation reflects stochastic de-repression or structured reorganisation of genome defence mechanisms^9,10^.

Sex differences in genome stability may further shape these responses. The ‘faulty male’ hypothesis proposes that the male germline is particularly vulnerable to genetic and epigenetic disruption due to the continuous and rapid production of sperm during spermatogenesis^11^. The increased exposure to DNA replication and meiotic cycles may amplify susceptibility to environmental stress, increasing the likelihood of failures in sRNA-mediated TE suppression and leading to disproportionate accumulation of mutations^9,12–15^. Understanding the interaction between epigenetic and genetic responses in the germ cells to environmental variation sheds light on their potential role in adaptation^7,16^. Recent evidence suggests that stress can preferentially reactivate evolutionarily young, potentially autonomous TE families, which are more likely to retain transposition capacity and thus pose a greater threat to genome integrity ^5^. This is particularly relevant in germ cells, where TE control relies on sRNA pathways such as the piRNA system and protein-folding buffers like HSP90, both of which are highly sensitive to physiological perturbation^17^. Disruption of these pathways under stress can weaken TE silencing, leading to increased TE activity and downstream effects on gene regulation^18, 19^. At the same time, sRNAs may contribute to adaptation by providing a rapid, reversible mechanism for regulating TE activity and gene expression in response to environmental change, complementing slower genetic evolution^20^.

Here, we combine experimental exposure to increased temperature in male and female zebrafish with DNA and RNA sequencing of the gonads to link temperature variation with changes in miRNA and piRNA profiles, TE activity and gene expression and functionally link them. We show that thermal stress drives a coordinated restructuring of the molecular genome defence. We identify a previously uncharacterised piRNA-TE-miRNA regulatory cascade that is maintained under heat stress in ovaries but collapses in testes. This collapse is associated with widespread TE mobilisation and genome instability, whereas ovaries preserve a highly connected regulatory network that safeguards germline integrity. Together, these findings define a multi-layer framework linking sRNA pathways, transposable elements, and gene regulation in germline stress responses, revealing a fundamental sex-specific divergence in how genome defence is maintained under environmental change.

## Results & Discussion

### Sex-specific transcriptional rewiring links stress response to sRNA regulation

Thermal stress induced distinct transcriptional responses in male and female gonads, with 378 differentially expressed genes (DEGs) in ovaries (210 upregulated, 168 downregulated; Fig. 1A,C) and 340 in testes (87 upregulated, 253 downregulated; Fig. 1B). A subset of shared DEGs, including *hspa8b, hsp90aa1.2, hsp70.3,* and *marcksl1a*, represent key nodes linking stress response pathways to RNA biogenesis and TE control^11,19,21^, and are predicted targets of differentially expressed miRNAs (Fig. 1D–F), directly connecting transcriptional and post-transcriptional regulation.

**Figure 1.**
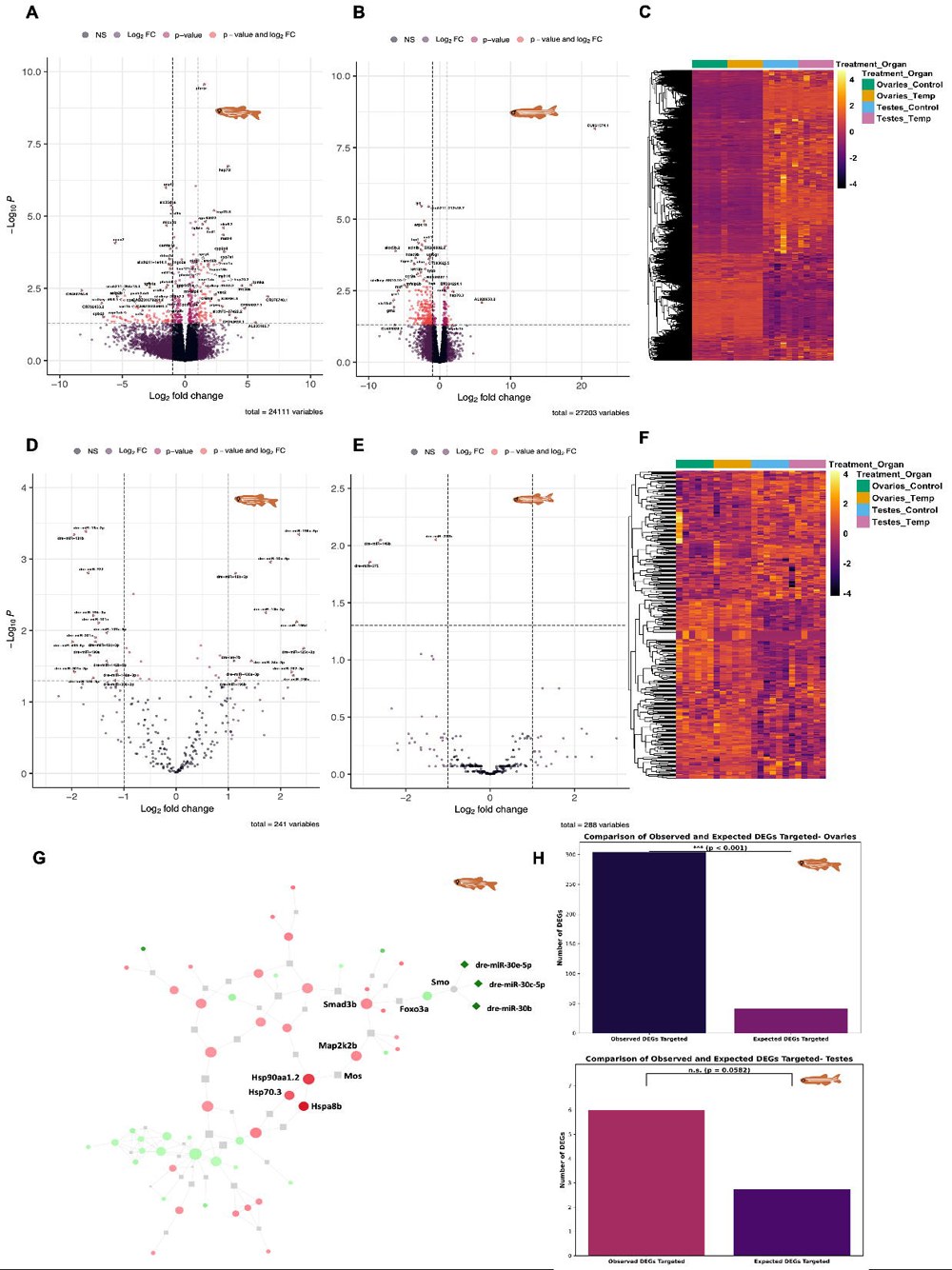
Differential expression and target analysis of genes and mature miRNAs in thermally stressed gonads. Differential expression analysis of RNA-seq data revealed strong gene upregulation and downregulation in female gonads **(A)** and a bias toward gene downregulation in male gonads **(B)** following thermal stress. Nine genes were significantly differentially expressed in both: *hspa8b, hsp90aa1.2, si:dkey-46i9.1, lgals9l3, pogzb, efh2, map2k2b, hsp70.3, and marcksl1a.* **(C)** Heatmap of normalised gene counts (log□CPM) from the DESeq2 output (design = ∼Treatment + Organ). **(D)** miRNA differential expression analysis revealed both up- and downregulation in female gonads **(E)** but predominant downregulation in male gonads. **(F)** Heatmap of normalized miRNA counts (log□CPM) from the DESeq2 output (design = ∼Treatment + Organ). Z-score-normalised log^CPM^ confirmed a congruent sample structure. **(G)** Multiomic clustering of significantly differentially expressed genes and miRNAs in ovaries via the OmicsNet minimal model. Circles = genes, squares = proteins, diamonds = miRNAs; red = upregulated, green = downregulated. Reduced *miR-30* cluster expression may predict gene upregulation. **(H)** MiRanda target prediction revealed significant enrichment of miRNA-gene interactions in thermally stressed ovaries but not in testes because of limited miRNA changes.

In ovaries, enrichment of proteostasis pathways (Supplementary Fig. 2A,C) coincided with miRNA-mediated release of repression on heat-shock genes (Fig. 1G), consistent with coordinated activation of chaperone systems that stabilise piRNA processing and TE silencing^19,22^. In contrast, testes exhibited enrichment of immune and inflammatory pathways (Supplementary Fig. 2B,D), a transcriptional signature associated with TE activation and chromatin destabilisation^16,23,24^, indicating a breakdown of genome defence mechanisms. PCA revealed tissue-specific expression shifts under thermal stress in both ovaries and testes (Supplementary Fig. 1). Hierarchical clustering showed coordinated covariation between miRNAs, piRNAs, transposable elements, and genes in ovaries, consistent with an integrated regulatory response, whereas testes displayed weaker coupling across these components. Together, this suggests a coordinated, protective response in ovaries but a more destabilised regulatory state in testes, linking gene expression changes to divergence in sRNA-mediated genome defence.

### Coupled disruption of miRNA and piRNA networks drives TE activation in testes

SRNA profiling revealed a coordinated regulatory axis linking miRNA-mediated gene control with piRNA-dependent TE repression. In ovaries, thermal stress induced extensive miRNA remodelling (41 differentially expressed miRNAs; Fig. 1D) alongside balanced piRNA expression (Fig. 2A,C), with significant enrichment of miRNA targets among differentially expressed genes (Fig. 1H). This indicates strong integration between transcriptional and post-transcriptional regulation, consistent with coordinated maintenance of genome stability.

**Figure 2.**
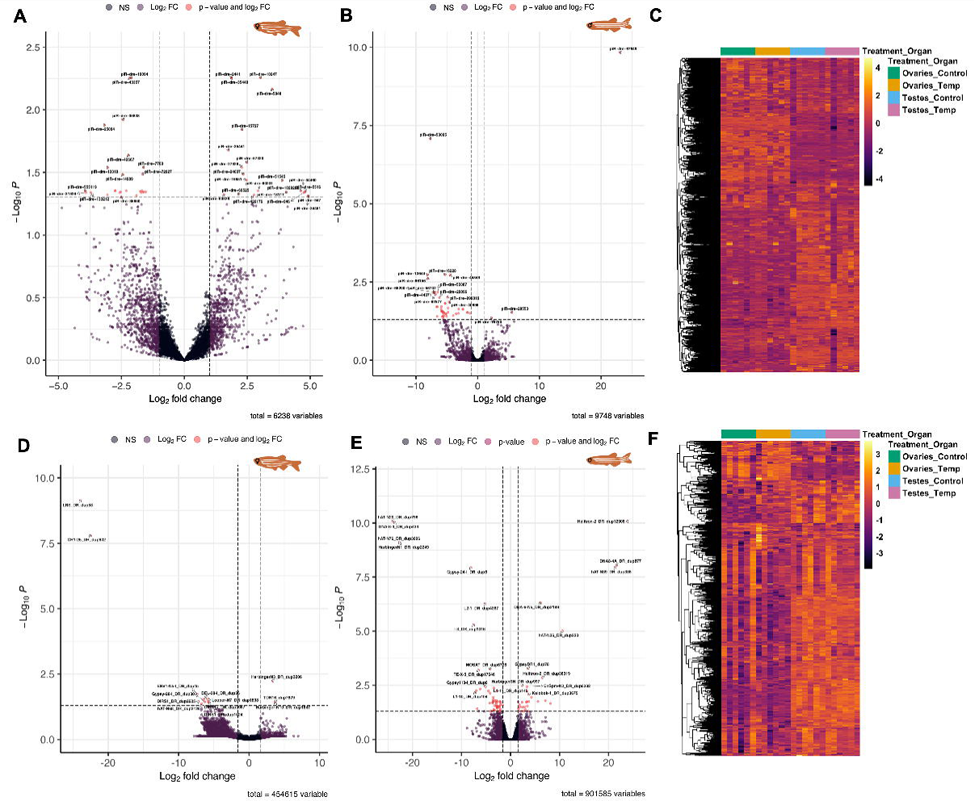
PiRNA and TE differential expression analysis from RNA-seq data. **(A-B)** Differential expression of piRNAs in thermally stressed gonads revealed enrichment and downregulation of piRNAs in female thermally stressed gonads and a strong bias for downregulation of the expression of piRNAs in male thermally stressed gonads. **(C)** Heatmap of normalised counts (log^2^ CPM) from the DESeq2 output of sRNA-seq data on piRNAs across all samples, with design = ∼ Treatment + Organ in dds object generation. **(D-E)** Differential expression analysis of TEs assayed in Telescope at the species level. **(F)** Heatmap of normalised counts (log^2^ CPM) from the DESeq2 output of sRNA-seq data on TEs across all samples, with design = ∼ Treatment + Organ in dds object generation.

In contrast, testes exhibited concurrent depletion of both miRNAs and piRNAs (Fig. 1D; Fig. 2B,F), indicative of a systemic failure of sRNA pathways^13,24^. This dual loss disrupts transcriptional buffering (via miRNAs) and TE silencing (via piRNAs), directly linking reduced sRNA abundance to increased TE activity. Depletion of structural RNAs in testes (Suppl. Fig. 4) further suggests impairment of RNA-mediated inheritance pathways^25,26^. Together with known roles of sperm-borne tRNA fragments in paternal transmission^27,28^, these changes point to potential inter- and transgenerational consequences.

Comparison of sRNA classes revealed pronounced asymmetry between males and females. Ovaries exhibited extensive remodelling of both precursor and mature miRNAs, including coordinated regulation of stress-responsive miRNA clusters, whereas testes showed minimal changes. Notably, shared miRNAs such as *dre-miR-462* and *dre-miR-731* (Suppl. Fig. 3A-C) displayed opposing regulation between sexes, suggesting divergent regulatory responses to thermal stress. Functional enrichment analysis indicated that ovarian miRNAs target pathways associated with cellular homeostasis and signalling, whereas testicular miRNAs were linked to neural and stress-related pathways (Suppl. Fig. 2).

Integration of miRNA target predictions with gene expression data further revealed coordinated regulation in ovaries (Fig.1, Suppl. Fig. 2), where depletion of *miR-30* family members coincided with activation of heat-shock pathways, consistent with release of translational repression. In contrast, testicular miRNA targets were not significantly enriched among differentially expressed genes, reflecting a breakdown of post-transcriptional regulatory control. Only three mature miRNAs were significantly altered in testes, all downregulated, reinforcing the limited capacity for miRNA-mediated regulation in this tissue. These findings further support that thermal stress preserves an integrated sRNA regulatory network in ovaries but induces a coordinated collapse of miRNA and piRNA pathways in testes, which is associated with increased TE activation.

### TE mobilisation reflects failure of sRNA-mediated genome defence

To determine the chromatin context of TE insertions, ATAC-seq signal profiles were analysed alongside RetroSeq-derived insertion sites. *De novo* insertions in heat-stressed testes frequently overlapped regions of elevated chromatin accessibility (Fig. 3D-red tracks indicate regions of open chromatin), consistent with integration into open and potentially regulatory genomic regions. However, this association was not absolute, as insertions were also detected in less accessible regions, indicating that TE mobilisation is not strictly confined to canonical open chromatin environments. The depletion of insertions observed on chromosome 4 (Fig. 3A–D) corresponded to regions of reduced ATAC-seq signal, suggesting that higher-order chromatin organisation on that chromosome contributes to spatial bias in TE integration. These findings support a model in which chromatin accessibility modulates, but does not fully determine, insertion probability, linking genome architecture to stress-induced patterns of transposition. Differences in sRNA regulation translated directly into TE dynamics. In thermally stressed ovaries, TE activity remained tightly controlled, with limited differential expression (Fig. 2D; Supplementary Fig. 5) and only a modest increase in insertion events (28□°C: 678; 34□°C: 764; Fig. 3B,D), predominantly in accessible chromatin regions. In contrast, testes exhibited widespread TE activation (Fig. 2E) and a near threefold increase in *de novo* insertions (28□°C: 787; 34□°C: 2,339; Fig. 3A,C), indicating a profound loss of genome defence. Z-score normalised expression profiles confirmed robust and consistent treatment effects across replicates (Fig. 2C,F). Insertion patterns further reflected this divergence. In ovaries, insertions were enriched in promoters and 3′UTRs (Fig. 3E–G), suggesting buffered regulatory effects, whereas in testes, insertions accumulated predominantly in intronic regions, consistent with active transposition in a permissive chromatin environment. These patterns demonstrate that loss of sRNA-mediated repression is associated with increased TE mobilisation in the male germline.

**Figure 3.**
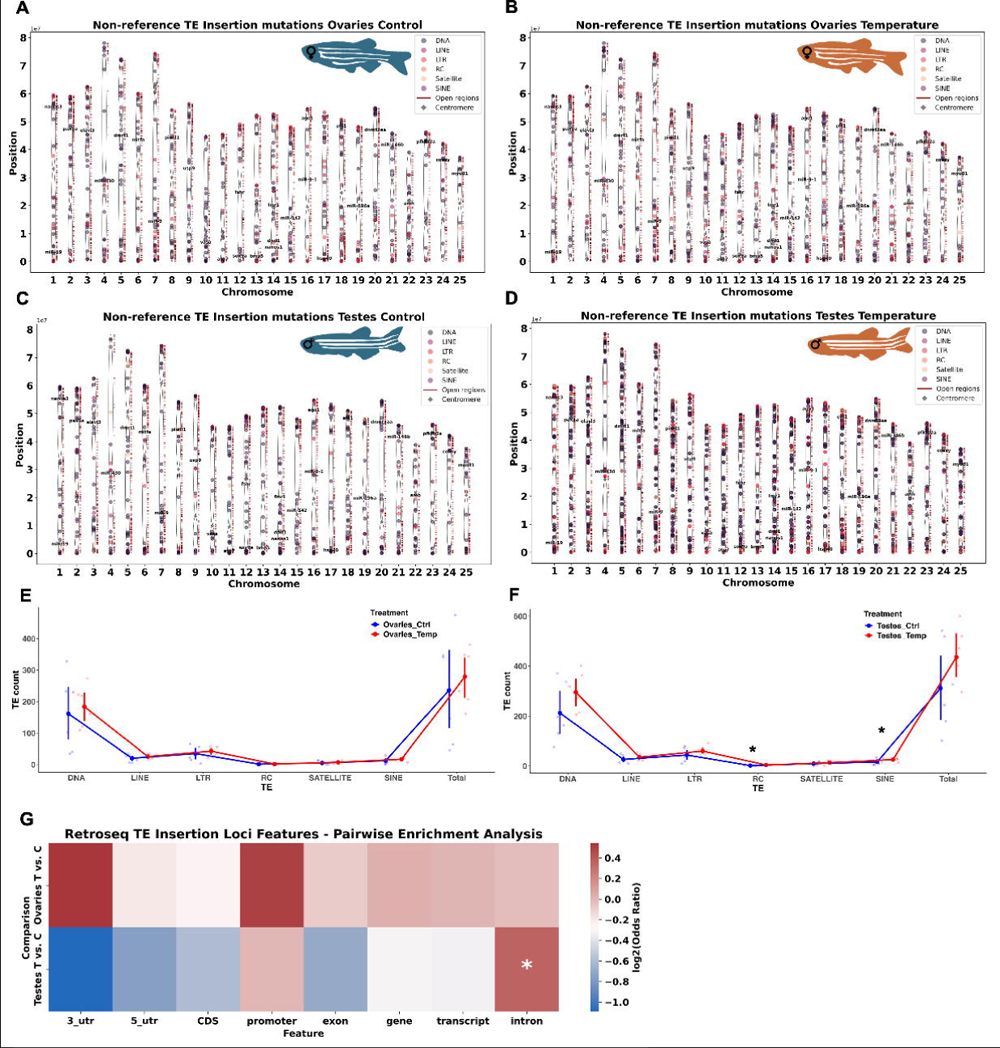
Nonreference transposable element insertions in the zebrafish genome as assayed via Retroseq. **(A-B)** Nonreference genome insertions caused by transposable elements in control versus temperature-stressed female gonads. **(C-D)** Nonreference genome insertions caused by transposable elements in control versus temperature-stressed male gonads. **(E-F)** Abundance of TE nonreference insertion mutations by family, tested for significance with GLM: rep_m1 <-glmer(Total ∼ Treatment + (1 | MaleID), family = Poisson, data = df), asterisks denote (*p* < 0.05). **(G)** Heatmap of log2-transformed odds ratios from Fisher’s exact tests showing enrichment of genomic features for TE insertions after thermal stress in ovaries (top row) and testes (bottom row). The colour intensity follows a red–blue palette, with red indicating greater enrichment and blue indicating lower enrichment. Asterisks mark features significantly enriched after Benjamini–Hochberg false discovery rate correction (*p* < 0.05).

Ovaries maintained TE repression, with limited expression changes (Fig. 2D), and modest insertion increases (Fig. 3B,D), consistent with intact miRNA-piRNA coordination. In contrast, testes exhibited widespread TE activation (Fig. 2E) and a near 3-fold rise in *de novo* insertions (Fig. 3A,C), particularly among young TE families. This pattern is consistent with known sex-biased piRNA efficacy ^13^ and the documented sensitivity of piRNA biogenesis to stress conditions ^19,24^. Insertion patterns reinforce these links: ovarian insertions occurred mainly in promoters and 3’ UTRs (Fig. 3E-G), suggesting buffered regulatory effects, whereas testicular insertions accumulated in introns, consistent with active transposition in a permissive chromatin environment. Thus, reduced piRNA output, combined with miRNA depletion, directly corresponds to elevated TE mobilisation in testes ^9,17^.

Thermal stress led to a global reduction in piRNA abundance in males (Fig. 2B-C); however, the ping□pong amplification signature remains relatively unchanged (Suppl. Fig. 5J). This indicates that the amplification cycle itself is not disrupted but instead operates on a reduced substrate pool. As the ping□pong signature reflects relative enrichment rather than absolute piRNA levels, its preservation suggests that mechanistic amplification efficiency is maintained, even as total piRNA output declines in thermally stressed testes. Consequently, reduced piRNA abundance limits the overall capacity of the pathway, resulting in insufficient production of secondary piRNAs to effectively repress TEs. These results reveal that the limiting factor in TE silencing is not the loss of amplification capacity, but a reduction in piRNA supply, placing the system below the threshold required for effective genome defence.

Kimura divergence-based repeat landscapes were used to infer the evolutionary age distribution of transposable elements (TEs), enabling resolution of recent versus ancient TE activity^29^. These analyses revealed modest, non-significant global shifts in TE divergence under thermal stress, with recent activity primarily attributable to LTR retrotransposons (Supplementary Fig. 6). To assess whether stress-induced TE activation preferentially involves evolutionarily young elements, we examined the divergence profiles of significantly differentially expressed TE families identified by Telescope. In thermally stressed testes, upregulated TE families were consistently enriched for low-divergence (recent) elements, particularly among DNA, rolling-circle (RC), satellite, and SINE families. In contrast, thermally stressed ovaries exhibited substantially lower TE activity overall, with no comparable enrichment of young TE families.

This bias towards low-divergence elements in testes shows that recently active, and potentially still-autonomous, TE families are preferentially mobilised under thermal stress, in line with models of stress-induced reactivation of epigenetically suppressed TEs^7,8,16,17^. While we did not perform a formal genome-wide enrichment test, the observed shift towards younger TE families provides strong indicative support for selective activation of evolutionarily recent elements. This indicates that stress does not activate TEs uniformly, but preferentially targets evolutionarily young elements that retain mobilisation capacity, revealing latent genomic instability.

At the genomic level, 225 significantly differentially expressed genes and TE families exhibited overlapping loci in thermally stressed ovaries, including 194 DNA, 9 LINE, and 22 LTR-associated loci, whereas no such overlaps were detected in testes (Suppl. Fig. 6F). These overlaps were notably absent from the long arm of chromosome 4, consistent with the reduced TE activity previously observed in this region. Genes overlapping TE loci were associated with developmental processes such as skeletal muscle development and neural tube closure, although these terms did not reach statistical significance.

Collectively, our results demonstrate a pronounced male-biased activation of evolutionarily recent TE families under thermal stress, with LTR retrotransposons representing a particularly high-risk class due to their capacity to alter regulatory and chromatin architecture^1^. The absence of insertions on the long arm of chromosome 4 further suggests that genomic organisation may impose regional constraints on TE mobilisation. Together, these findings support a mechanistic model in which loss of miRNA-piRNA coordination in testes leads to reduced piRNA availability, failure of TE repression, and selective activation of young, mobile elements, resulting in widespread genome instability under thermal stress.

### TE-gene interactions and selection signals couple genomic instablity to functional outcomes

TE insertions intersected functionally relevant loci, including germline regulators such as *nanos3, dnd1*, and the *dre-miR-430* loci (Fig. 3), linking TE mobilisation directly to pathways controlling germline development and early embryogenesis^30–32^. Given the broad regulatory scope of *miR-430*^25^, disruption of these loci integrates TE dynamics with sRNA-mediated gene regulation.

Genome-wide analyses further revealed signatures of balancing selection (Fig. 4), with elevated Tajima’s D indicating selection on standing variation^33,4,18,19,22,34–39^. Genome scans revealed sex-specific signals of differentiation (Fig. 4), with elevated Tajima’s D indicating selection on standing variation^33^. Testes carried a higher burden of regulatory variants (Suppl. Fig. 7) that overlapped with regions of TE activity and sRNA disruption, suggesting feedback between TE mobilisation, regulatory variation, and gene expression. Together, these patterns support a model in which genetic variation and epigenetic disruption converge under thermal stress, with TE activation acting as both a source of genome instability and a potential contributor to adaptive variation^4,8,16,17,22,34–39^.

**Figure 4.**
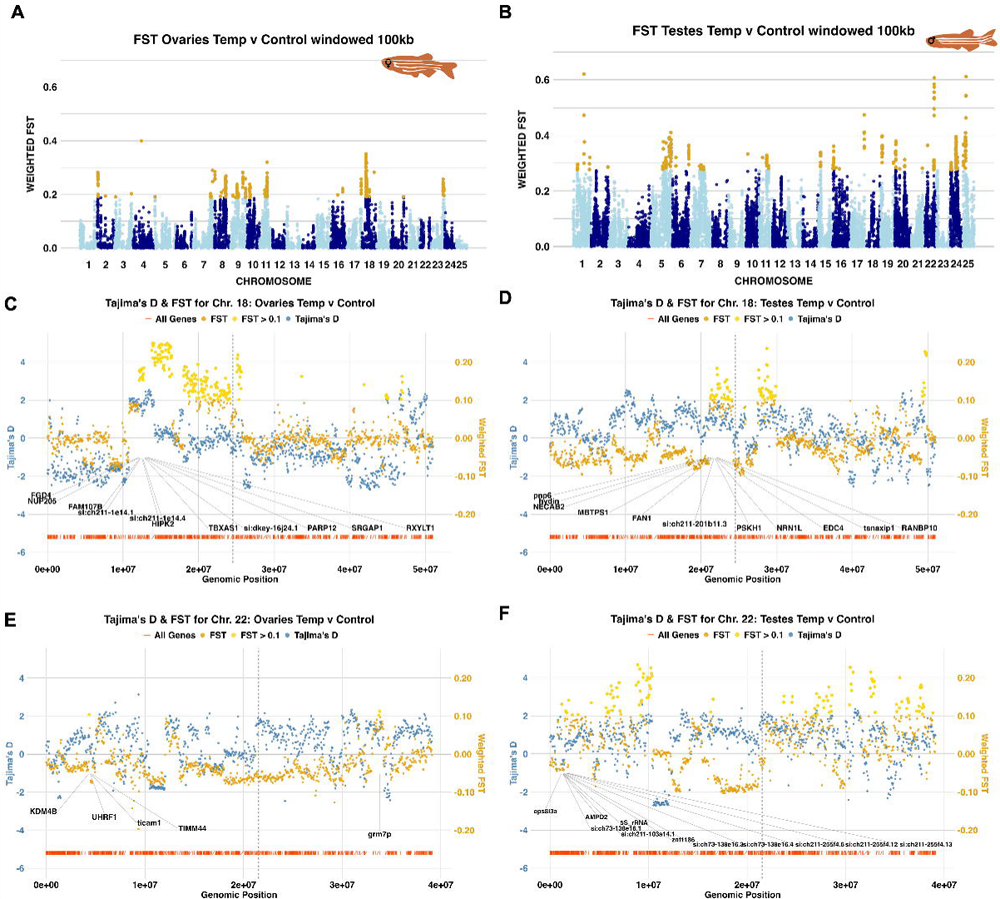
FST and Tajima’s D analysis of the genomic DNA-seq data. **(A-B)** Genome-wide weighted F_ST_ analysis (window size 100 kb), *p* = < 0.05; points in gold are in the 99^th^ percentile. **(C-F)** Chromosomal-level view of FST and Tajima’s D statistics, with gene loci annotated in vertical red bars. Genes under areas of FST peaks > 0.1 are annotated. The centromere location is annotated by a vertical dashed grey line. Chromosome 18 for ovaries **(C)** and testes **(D)** following thermal stress. Chromosome 22 for ovaries **(E)** and testes **(F)**.

### Transgenerational consequences reinforce functional integration

Thermal stress effects extended beyond the exposed generation. Parental heat exposure altered offspring gene expression (Suppl. Fig. 9A-B), and reduced reproductive success, with a strong paternal effect on embryo mortality (Suppl. Fig. 9C-D). Analysis of embryo survival revealed a significant paternal effect, paternal thermal stress was consistent with a significant increase in 24-hour embryo mortality (binomial GLM: *p□*=□0.018), regardless of the mother’s treatment (Suppl. Fig. 9D). These outcomes link disrupted sRNA pathways and TE activity in testes to functional consequences for fertility and inheritance^20,40^. These findings suggest a link between stress-induced disruption of germline genome defence and reduced reproductive fitness, indicating that sRNA-mediated regulation may be important for maintaining both genome stability and heritable integrity under environmental stress.

### Multi-omic regulatory network analysis

To resolve how thermal stress affects regulatory interactions across the epigenetic-genetic axis, we reconstructed integrated networks linking miRNAs, piRNAs, transposable elements (TEs), and gene expression. Networks were constructed by integrating experimentally supported interactions, including miRNA-gene, piRNA-TE, and TE-gene relationships, with differentially expressed features, and were visualised in Cytoscape (see interactive networks in Supplementary Files 16–17). In testes, the resulting network was minimal, comprising only two miRNAs (*dre-miR-146b* and *dre-miR-222b*) and two target genes (*hsp70.3* and *hspa8b*), with depleted miRNAs corresponding to upregulated target genes, consistent with canonical miRNA-mediated repression (Suppl. File. 16). In contrast, ovaries exhibited a highly connected network spanning all molecular levels (Supplementary File 17). A prominent subnetwork centred on the DNA transposon *DNA□3*□*1_DR*, which formed multiple interactions with both piRNAs and gene targets. This TE was linked to a gene regulatory module involving *acsl1a*, an upregulated gene under thermal stress, predicted to be targeted by multiple downregulated miRNAs (including *dre-miR-107a-5p*, *dre-miR-101a* and members of the *miR-19* family). This configuration indicates coordinated release of miRNA-mediated repression alongside TE-associated regulatory interactions. DNA□3□1*_DR* was connected to a gene regulatory module via *acsl1a*, a gene that was upregulated in this condition. The gene *ascl1a* is an ortholog to the human *ASCL1* gene and plays key roles in neurogenesis^41^. The gene *acsl1a* was associated with DNA□3□1*_DR* and was predicted to be regulated by a set of downregulated miRNAs, including *dre-miR-107a-5p, dre-miR-101a, dre-miR-19a-3p, dre-miR-190a, dre-miR-19b-3p, dre-miR-101b*, and *dre-miR-19c-3p*. This configuration supports a model in which the simultaneous reduction of miRNA-mediated repression and potential TE-associated regulatory effects influence gene expression.

The TE linked to this gene, *DNA□3□1_DR*, belongs to the PIF-Harbinger family of Class II DNA transposons, and its integration across both piRNA-TE and TE-gene interaction layers highlight a previously underappreciated role for DNA transposons as intermediates in regulatory networks. The presence of TE-centred modules uniquely in ovarian networks supports a model in which transposable elements facilitate the coordination of sRNA-mediated gene regulation under stress^2^. Together, these findings suggest that transposable elements may be components of regulatory networks linking sRNAs to gene expression. We define a multi-layer regulatory architecture revealing a striking divergence in regulatory architecture: thermally stressed ovaries retained a highly connected, multi-layer network, whereas testes exhibited a near-complete collapse of regulatory connectivity. This contrast indicates that thermal stress does not simply alter gene expression but selectively disrupts higher-order regulatory organisation in the male germline. Consistent with this, no transcription factor-driven regulatory structure was detected, suggesting that sRNA pathways represent the dominant drivers of network organisation under stress.

## Conclusions

We identified pronounced sex differences in the molecular response to thermal stress in gonads, where ovaries preserve an integrated, multi-layer genome defence network, while testes undergo a coordinated collapse of sRNA-mediated regulation, resulting in widespread TE activation. This divergence supports the ‘faulty male’ hypothesis and reflects fundamentally different strategies of genome maintenance, with ovaries maintaining robust miRNA-piRNA coordination, sustained TE repression, and stable transcriptional regulation, whereas testes exhibit reduced TE surveillance, immune-associated transcriptional signatures, and disruption of RNA-mediated inheritance pathways. TE insertions at key fertility-associated loci (*nanos3, dnd1,* and *dre-miR-430*) and altered structural RNA profiles in testes are associated with changes in germline function and offspring fitness^42^. In parallel, signals of selection within genic regions suggest that thermal stress interacts with standing genetic variation, particularly in pathways related to sRNA function and TE control, highlighting a potential interface between genome instability and adaptive potential. Collectively, our results support a model in which thermal stress exposes sex-specific vulnerabilities in germline genome defence, characterised by maintenance of an integrated piRNA–TE–miRNA regulatory network in ovaries and its collapse in testes. Our study establishes that transposable elements are not solely targets of genome defence, but active components of multi-layer regulatory systems linking sRNAs to gene expression. The disruption of this cascade under environmental stress provides a mechanistic framework for understanding how genome stability, fertility, and heritable variation are shaped in a changing environment.

## Methods

### Zebrafish husbandry

We used adult (1 year old) male and female zebrafish from the AB wild-type strain obtained from the Zebrafish International Research Centre (ZIRC, Oregon, USA) and reared and maintained at the Controlled Environment Facility, University of East Anglia, UK. The fish were raised to sexual maturity and kept under standard laboratory conditions at a temperature of 27°C with a 14 h:10 h light:dark cycle. The fish were fed *ad libitum* three times per day with a mixture of live *Artemia* (ZM Systems, UK) and dry food Sparos (Medium granular, ZM Systems, UK). Water quality was monitored with a JBL Pro Aqua Test Freshwater Test Kit (JBL, Germany). The light intensity was monitored with a Multicomp PRO MP780118 digital light meter (MulticompPro/Farnell, UK). The water pH was assessed with a pH meter (Fisherbrand Accumerator AET15 pH tester; Thermo Fisher, USA). The results can be found in Suppl. Fig. 8 to display the temperature, light, hardness, carbon dioxide and oxygen-related parameters.

### Experimental setup

For the experiments, zebrafish were housed in groups of 12 fish per 3 L tank at equal sex ratios (six males and six females), and the tanks were either maintained at 28°C (control temperature) or 34°C (high temperature). The fish were exposed to the experimental treatment for two weeks and then euthanised via the Schedule 1 method under the Home Office Project Licence P0C37E901. Gonads were collected by dissection on ice, homogenised in a sterile dish to produce two aliquots of each gonad, frozen in liquid nitrogen and stored at −80°C until further processing. Gonads were collected across two blocks from above set-up, with 6 biological samples per following groups: ovaries control, ovaries temperature, testes control, testes temperature. For transgenerational analysis fish were then bred in pairs (1:1 equal sex ratios) following heat stress treatments. Adult fish were crossed in the following: 28 °C F x 28 °C M, 28 °C F x 34 °C M, 34 °C F x 28 °C M and 34 °C F x 34 °C M, to profile for maternal or paternal impacts and inhertance. Offspring were reared at 28 °C before being frozen in pools of 10 24 h.p.f embryos in liquid nitrogen.

### DNA and RNA extraction

DNA was extracted from one aliquot of each homogenised gonad via Qiagen DNeasy Blood and Tissue kits (Cat. No./ID: 69504) according to the manufacturer’s instructions, but we omitted the RNase step. For RNA extraction, we used the Zymo Quick-RNA miniprep plus kit (R1057) according to the manufacturer’s instructions but used Zymo-Spin IC columns (C1004□50) because of the low amount of tissue available (< 5 mg).

### Quantitative PCR

To perform qRT-PCR, 200 ng of total RNA was isolated from pooled embryos (10 x 24 h.p.f; frozen in liquid nitrogen) and used to generate cDNA (PCR biosystems UltraScript cDNA synthesis kit PB30.11-10). For the qRT-PCR reactions 2X syGreen mix (PCR biosystems PB20-15-06) to generate 20 µL reactions on a Bio-Rad CFX96 real-time detection system. Primers were diluted to final concentration of 10 µM in each reaction. The thermal cycling protocol consisted of 50□°C for 2 min, 95□°C for 10□s, followed by 40cycles of 95□°C for 15□s and 60□°C for 60□s. A final denaturation at 95□°C for 15□s and an annealing/extension step at 60□°C for 15□s were performed prior to melt curve analysis (60–95□°C, 0.5□°C increments). Gene expression was quantified using the ΔΔCt method, with Ct values normalised to GAPDH and relative expression calculated as 2^ΔΔCt. Primer sequences in 5’ -> 3’ orientation for GAPDH: Forward: GTGGAGTCTACTGGTGTCTTC, Reverse: GTGCAGGAGGCATTGCTTACA ^43^. HSP90 Forward: TTCGGTGTGGGCTTTTATTC, Reverse: TTAATGCGCTTTTCCTCCAC^44^. HSP70l Forward: TACGAGGGCATCGACTTCTA, Reverse: CAGTGCTTTCTCCACAGGAT^44^.

### Library preparation and whole-genome sequencing

DNA libraries were prepared via the LITE library Prep method (Earlham Institute, Norwich, UK, LITE v1.0) and sequenced on an Illumina NovaSeq 6000 S4 with 150 bp paired end reads to achieve coverage of 34 X or greater. The adapters used in LITE library preparation were [i7] and [i5] with 9 bp indices, where the sequences ranged from 5’-3’, and appeared in the sequencing data as the reverse complement (P7 in read 1 and P5 in read 2; P7: CAAGCAGAAGACGGCATACGAGAT[i7]GTCTCGTGGGCTCGGAGATGTGTATAAG AGACAG; P5: AATGATACGGCGACCACCGAGATCTACAC[i5]TCGTCGGCAGCGTCAGATGTGTA TAAGAGACAG).

### Library preparation and mRNA sequencing

The mRNA sequencing libraries were prepared via the New England Biolabs NEBNext Ultra II Directional RNA library prep (rRNA depleted library prep) to generate 150 bp paired-end reads at a depth of 20 million reads and a PhiX spine-in of 1%. The adapters used in the RNA-seq in the 5’ to 3’ orientation were as follows: forward: 5’-3’: AGATCGGAAGAGCACACGTCTGAACTCCAGTCAC; reverse: AGATCGGAAGAGCGTCGTGTAGGGAAAGAGTGTA. This service was provided by Source Bioscience, Cambridge, UK. Sequencing was performed on a NovaSeq 6000.

### Library preparation and sRNA sequencing

The sRNA sequencing libraries were prepared via the Qiagen QIASeq miRNA library Prep Kit (cat. No. 331502) to generate 100 bp paired end reads sequenced to a depth of 10 million reads. The adapters used in the sRNA-seq in the 5’ to 3’ orientation were as follows: forward: AACTGTAGGCACCATCAAT; reverse: GTTCAGAGTTCTACAGTCCGACGATC. A 1% PhiX spike was used. This service was provided by Source Bioscience, Cambridge, UK. Sequencing was performed on a NovaSeq 6000.

### Bioinformatic analyses

#### DNA transposable element analysis

The reads were trimmed with *fastp* version 0.23.2^45^. The reference zebrafish genome GRCz11 version 107 was obtained from Ensembl (Danio_rerio.GRCz11.dna.primary_assembly.fa.gz) and indexed with bwa version 0.7.17^46,47^. Reads were aligned to the reference genome, with the addition of read groups via BWA-MEM and samtools v1.11^48^. The code used was: bwa mem -t 4 -M -R “@RG\tID:$RGID\tSM:$SM\tPL:$PL\tLB:$LB\tPU:$PU” $ReferenceO $filea $fileb | samtools fixmate - - | samtools sort -T ${thisDirectory}bwamem/tmp_${ID} -O bam -o ${thisDirectory}bwamemL3/${outID}_ZFISH-aligned_sorted_RGs.bam. The sequencing lanes were then merged via picard v2.24.1, MergeSamFiles^49^.

#### Abundance of non-reference TE insertions

Retroseq was used to call nonreference TE insertions in the genome^50^. It was installed following the instructions (https://github.com/tk2/RetroSeq/blob/master/README.md) and run in a conda environment (the full list of packages and versions in this environment are listed in Appendix 9). The PopoolationTE2 preparation pipeline^51^ was used to generate the GRCz11.teseqs.fasta file, along with bedtools version 2.29.2^52^. The TE.bed file was obtained from the TEsmall database to be compatible with the GRCz11 v107 Ensembl reference genome ^53^. The resulting GRCz11.teseqs.fasta file was used to generate the reference TE sequence file for the Retroseq pipeline.

Retroseq was run in a conda environment with SAMtools 0.1.9, Bedtools 2.29.2 and Exonerate 2.2.0^54^ using default settings for the discovery and calling phases. The final VCF was filtered in three stages. First, calls that were unique for the experimental groups were filtered out and analysed, and any TE insertions in the control groups were thus removed from the experimental group vcf file via a unique.py script. Second, to reduce false positive insertion mutations, any TE insertion mutation calls within 100 bp of a known TE insertion were removed via the following steps: bedtools window -b knownTEs.bed -an input.vcf -v -w 100 -header >> output.vcf. This was done via BCFtools version 1.10.2, bedtools version 2.29.2 and VCFtools version 0.1.16 ^48,55^. Final filtering for plotting was conducted to select calls based on the INFO tags: FL=8 & GQ>750, to select the most confident calls via script filter.py. Plots were generated via the script retroseq_ultimate_phenogram_genes.py. To identify TE insertion mutations overlapping with a gene, we ran: bedtools intersect -an input.vcf -b genes.bed -wa -wb > output_with_genes.vcf. For the phenogram plots, the centromere locations were obtained from Table 2 in^56^. These locations were aligned against previous *danRer4* and *GRCz11* reference genomes via BLAST for validation. TE class plots were generated via re, pandas and matplotlib; script class_chart.py; and test.py for plotting. To assess the location of TE insertion mutations, heatmaps of ^log2-^transformed odds ratios from Fisher’s exact tests showing enrichment of genomic features with Benjamini–Hochberg false discovery rate correction (*p* < 0.05) was generated. To further analyse the impact of the TE insertion mutations discovered with Retroseq, these vcf files were used and run with Ensembl VEP^57^ online to profile broader impacts and consequences of TE insertions. FishTEA scripts were generated and used to intersect differentially expressed gene and TE loci^58^.

### Genomic divergence analysis

The sequencing reads were initially trimmed with Fastp v0.23.2 to remove adapters^45^. The reference genome used in this project was *Danio rerio* GRCz11, which was accessed from Ensembl, release 107 (https://ftp.ensembl.org/pub/release-107/fasta/danio_rerio/dna/) file “Danio_rerio.GRCz11.dna.primary_assembly.fa.gz”,^59^. The reference genome was indexed with BWA v0.7.17^46^. To align our reads with the reference genome and attach read groups bwa mem and samtools v1.11, *samtools fixmate - -* and *samtools sort -T* were used to sort the reads and generate bam files^46,48^. The samples from multiple lanes were merged with Picard *MergeSamFiles*^49^. The read groups were checked with SAMtools view and SAMtools addreplacerg. Bam files were processed with *samtools view -f 0×2 “${BAM}” -b -o “${BAM}”_out.bam* to remove unpaired reads. Picard was used to mark duplicates^49^. SAMtools *flagstat* and samtools *depth* were used to generate mapping statistics^48^. The generated bamfiles were indexed with *the SAMtools index* to split variant calling into individual chromosomes for increased processing speeds^48^. To call variants, BCFtools was used with the following script: *BCFtools mpileup -a AD,ADF,ADR - -q 30 -Q 30 -f Danio_rerio.GRCz11.dna.primary_assembly.fa -b bam.list | BCFtools call -mv -f GQ | BCFtools filter -g 3 -G 3 -e ‘DP < 85 || DP > 1600 || F_MISSING > 0.5 || QUAL < 30’ -Oz - o ${thisDirectory}bcf/${sample_name}_bcftools_zf.vcf.gz*^48^. DP parameters were set to 1/6 (1/3×1/2) of the total summed average coverage per sample. The output files were then merged with *BCFtools concat*^48^.

### Variant analyses

To analyse variants, SnpEff^60^ and SnpSift^61^ were used to generate annotated vcf files. These were filtered by significance by filtering CC_TREND (Cochrane□Armitage trend modelling), *p* = < 0.001 with SnpSift caseControl, and by snp impact, e.g., high. To generate eigenvec and eigenval files, PLINK v2.0.0 was used^62^. VCFtools was also used to conduct FST and Tajima’s D analyses^55^.

### RNA-seq and sRNA-seq analysis

The NF-core RNA-seq pipeline version 1.4.2 was used to analyse the RNA-seq data (https://nf-co.re/rnaseq/1.4.2/). The full bash scripting can be found in the appendices below. *The* GRCz11 version 107 reference genome was used as above, along with the accompanying version 107 Danio_rerio.GRCz11.107.gtf.gz file. sRNA-seq analysis of miRNAs (hairpin and mature) was performed via another NF-core pipeline: smRNA-seq version 2.0.0 (https://nf-co.re/smrnaseq/2.0.0)^63^. All the submission scripts and configuration files can be found in Appendices 1--5. MiRNA mature and hairpin fasta files were accessed from miRbase in September 2022 https://mirbase.org/download/^64^. Danio_rerio.GRCz11.107.gtf was accessed from the Ensembl database at https://ftp.ensembl.org/pub/release-107/gtf/danio_rerio/.

### Miranda miRNA analysis

For sensitivity and statistical confidence of miRNA□target gene associations in the 3’ untranslated region (UTR) of mRNAs by miRNAs, the miRanda version 3.3a pipeline was used with default settings^65^. MiRanda was run with a fasta file of mature zebrafish miRNAs and 3’ UTRs to scan and generate scored putative matches in the output. Full scripts and files can be accessed here: https://github.com/alicegodden/paternalsocstress/tree/main/miRanda. Zebrafish UTRs were obtained from TargetScan^66^, and the mature miRNA fasta file was obtained from miRbase^64^. Multiomic network integration was performed via OmicsNet (https://www.omicsnet.ca)^67^, which applies the minimal interaction model to link differentially expressed genes and miRNAs, enabling visualization of regulatory relationships and pathway-level connectivity.

### GO term analyses

To perform GO term analyses of the miRNAs, the predicted target genes of the miRNAs of interest were taken from the miRanda analysis. All significantly predicted target genes were subjected to GO term analyses. The RNA-seq dataset was used as a background list, which included all genes with a *p* value following DESeq2 analysis. ShinyGO version 0.80 (http://bioinformatics.sdstate.edu/go),^68^, was used to perform all the GO term analyses. GO term enrichment was performed via hypergeometric distribution with FDR correction and is included in the plots. The results were plotted with custom python scripts.

### Transposable element, structural RNA and piRNA analyses

TEtranscripts v2.2.3 was used for RNA-seq data to identify TEs that were differentially expressed at the family level^69^. The TETranscripts pipeline was run using -g GRCz11 genome settings. The strandedness of our RNA-seq data was checked with the MultiQC report^70^. TETranscripts were executed with the following settings: sorted_bam_files.bam --GTF Danio_rerio.GRCz11.107.gtf --TE GRCz11_Ensembl_rmsk_TE.gtf --mode multi --project Ovaries_Temp_vs_Control --minread 1 -i 100 --padj 0.05 --outdir results. The GRCz11_Ensembl_rmsk_TE.gtf annotation file was obtained from (https://labshare.cshl.edu/shares/mhammelllab/www-data/TEtranscripts/TE_GTF/).

Telescope v1.0.3 was run with the RNA-seq data to obtain species-level differentially expressed TEs ^71^ as follows: telescope assign OC1.markdup.sorted.bam_sorted.bam GRCz11_Ensembl_rmsk_TE.gtf --attribute transcript_id. To find and assign TE family and class information, the Dfam database was used https://www.dfam.org/home^72^. To create a raw count matrix for DESeq2 analysis for the TEsmall and Telescope results, the R script in Appendix 14 was used. TEsmall version 2.0.2 was run using the sRNA-seq data ^53^. TEsmall was executed with: TEsmall -a AACTGTAGGCACCATCAAT -g GRCz11 -f raw.fastq.gz -l 4MT -p 60. For ping-pong signature analysis sRNA sequencing reads were processed using the TEsmall pipeline and aligned to the zebrafish reference genome (GRCz11) using Bowtie (v1.3.1) with parameters allowing only perfect matches (−v 0) while retaining multi-mapping reads (−a −m 100). Reads corresponding to ribosomal RNAs were removed prior to alignment. piRNA-sized reads (24–32 nt) were retained for downstream analysis. Ping□pong amplification was quantified using a custom Python script applied to sRNA BAM files. piRNA-sized reads (24–32 nt) were extracted, and strand-specific 5′ coordinates were defined (start for sense reads, end −1 for antisense). Overlaps between sense and antisense 5′ ends were calculated within a 0–30 nt window, and enrichment at ∼10 nt (observed at 9 nt due to alignment offset) was used to identify ping□pong signatures. Ping□pong strength was quantified as the ratio of overlap counts at 9 nt to the mean counts at all other positions, providing a normalized enrichment metric per sample. Statistical comparisons between groups were performed using SciPy.

FishPi is a novel piRNA target prediction tool for the prediction of complementary piRNA:TE binding^73^. FishPi can be accessed here: https://github.com/alicegodden/fishpi. FishPi uses the following packages and versions: python 3.11, Tkinter (included in python 3.11), Pillow 10.0.1 and Matplotlib 3.8.0.

RepeatMasker^74^ version 4.1.1 was used to profile TE annotation, TE divergence and age (Kimura substitution estimated from DNA-seq data to analyse TE activity in the genome. *De novo* assemblies were generated by trimming raw reads with Fastp and concatenating lanes together. Megahit was used to generate assemblies per default settings for the DNA-seq paired-end data^75^.

Repeatmasker was run with the following settings: *RepeatMasker -species zebrafish -par 5 - s -a -nolow -no_is*. Repeat landscapes were generated with *Perl calcDivergenceFromAlign.pl -s file_divsum.* General linear modelling was used to conduct statistical analysis of the shift in genome expansion, with negative binomial modelling selected. The modelling results can be found in Supplementary Tables 1-2.

### Graphical plotting

RStudio version 4.1.1 ^76^, and the following packages were used in the analysis and presentation of the data in this project. These included: DESeq2 version 1.34.1^77^, Tidyverse version 2.0.0 ^78^, EnhancedVolcano version 1.12.0^79^, ggplot2 version 3.4.2^80^, biomaRt version 2.50.3^81^, org.Dr.eg.db version 3.14.0 ^82^, gplots version 3.1.3^83^, RColorBrewer version 1.1-3^84^ and Readr version 2.1.4^85^, ggVennDiagram version 1.2.3^86^, ggpubr version 0.6.0^87^, tidyr version 1.3.0^88^ and viridis version 0.6.4^89^, purrr version 1.0.2 ^90^, dplyr version 1.1.4^91^, qqman version 0.1.9^92^, qvalue version 2.34.0^93^, ggrepel version 0.9.4^94^, knitr version 1.45^95^, MASS version 7.3-60.0.1 ^96^, pscl version 1.5.9,^97^. To profile a top-level view of how the germline genome responded to thermal stress, differential expression lists for all sRNAs, TEs and genes were used. In addition, the results from FishPi (piRNA:TE), FishTEA (TE:gene) and miRanda (miRNA:gene) were combined to create nodes and edges. For multiomic network generation and integration Cytoscape v3.10.4^98^ was used to visualise nodes and edges. For all scripts used in this analysis, see https://github.com/alicegodden/thermalstress.

## Abbreviations

DE/DEG: differentially expressed/differentially expressed gene
miRNA: microRNA
piRNAs: piwi-interacting RNAs
sRNA: small RNA
TE: transposable element
tRNA: transfer RNA

## Declarations

### Ethics declarations

All zebrafish work was completed in accordance with institutional, local and national laws and governing bodies.

### Consent

*Not applicable*.

### Data availability

All sequencing data, including DNA-seq, RNA-seq and sRNA-seq data generated for this project, are available under accession number PRJEB72689 (under embargo until acceptance for publication) https://www.ebi.ac.uk/ena/browser/view/PRJEB72689. All scripts are available at the git repository: https://github.com/alicegodden/thermalstress, with supplemental data files here: https://github.com/alicegodden/thermalstress/blob/main/supp_files/README.md, and FishPi is published here^99^.

## Author contributions

The studies were conceptualized and designed by A.M.G. and S.I. Data analysis and visualization were carried out by A.M.G. J.C.D.C., R.S and C.D provided support with zebrafish handling and sampling. All the code was conceived and generated by A.M.G. Supervision and funding were provided by S.I. The first draft of the manuscript was written by A.M.G. and S.I. All the authors revised and approved the manuscript.

## Extended data

### Supplementary materials and methods

See the file “Godden et al_SupplMat.doc” for supplementary methods, figures, tables and legends.

### Supplemental files for data analysis

All files listed below are available here: https://github.com/alicegodden/thermalstress/blob/main/supp_files/README.md

1 – Water testing aquariums at 28°C and 34°C for 2 weeks of treatment – water testing.xlsx

2– DESeq2 analysis of differential gene expression – Testes: 2– te_tocontrol_deseq2.csv

3– DESeq2 analysis of differential gene expression – Ovaries: 3– ov_tocontrol_deseq2.csv

4– DESeq2 analysis of differential TEs (TETranscripts) expression– Testes: 4– te_tetrans_deseq2.csv

5– DESeq2 analysis of differential TE (TETranscripts) expression– Ovaries: 5-ov_tetrans_deseq2.csv

6– DESeq2 analysis of differential hairpin miRNA expression– Testes: 6-TESTES_DEDUP_HAIRPIN_TVsC_volcano.tsv

7– DESeq2 analysis of differential hairpin miRNA expression– Ovaries: 7-OVARIES_DEDUP_HAIRPIN_TVsC_volcano.tsv

8– DESeq2 analysis of differential mature miRNA expression– Testes: 8-TESTES_DEDUP_MATURE_TVsC_NA_use2.tsv

9– DESeq2 analysis of differential mature miRNA expression – Ovaries: 9-OVARIES_DEDUP_MATURE_TVsC_NA_use.tsv

10– Analysis of differential TE (telescope) expression – testes: 10-telescope_male_raw.csv.zip

11– Analysis of differential TE (Telescope) expression– Ovaries: 11-telescope_female_raw.csv.zip

12– Analysis of differential piRNA expression: 12-male_tesmall_pirna_volcano.csv

13– Analysis of differential piRNA expression – Ovaries: 13– female_tesmall_pirna_volcano.tsv

14– DESeq2 analysis of differential structural RNA expression– Testes: 14– male_structRNA_tesmall_deseq2.csv

15– DESeq2 analysis of differential structural RNA expression – Ovaries: 15– female_structRNA_tesmall_deseq2.csv

