## Supplementary Tables and Figures for "Sex differences in sRNA mediated transposon regulation under thermal stress in zebrafish germ cells"

### Slide 1
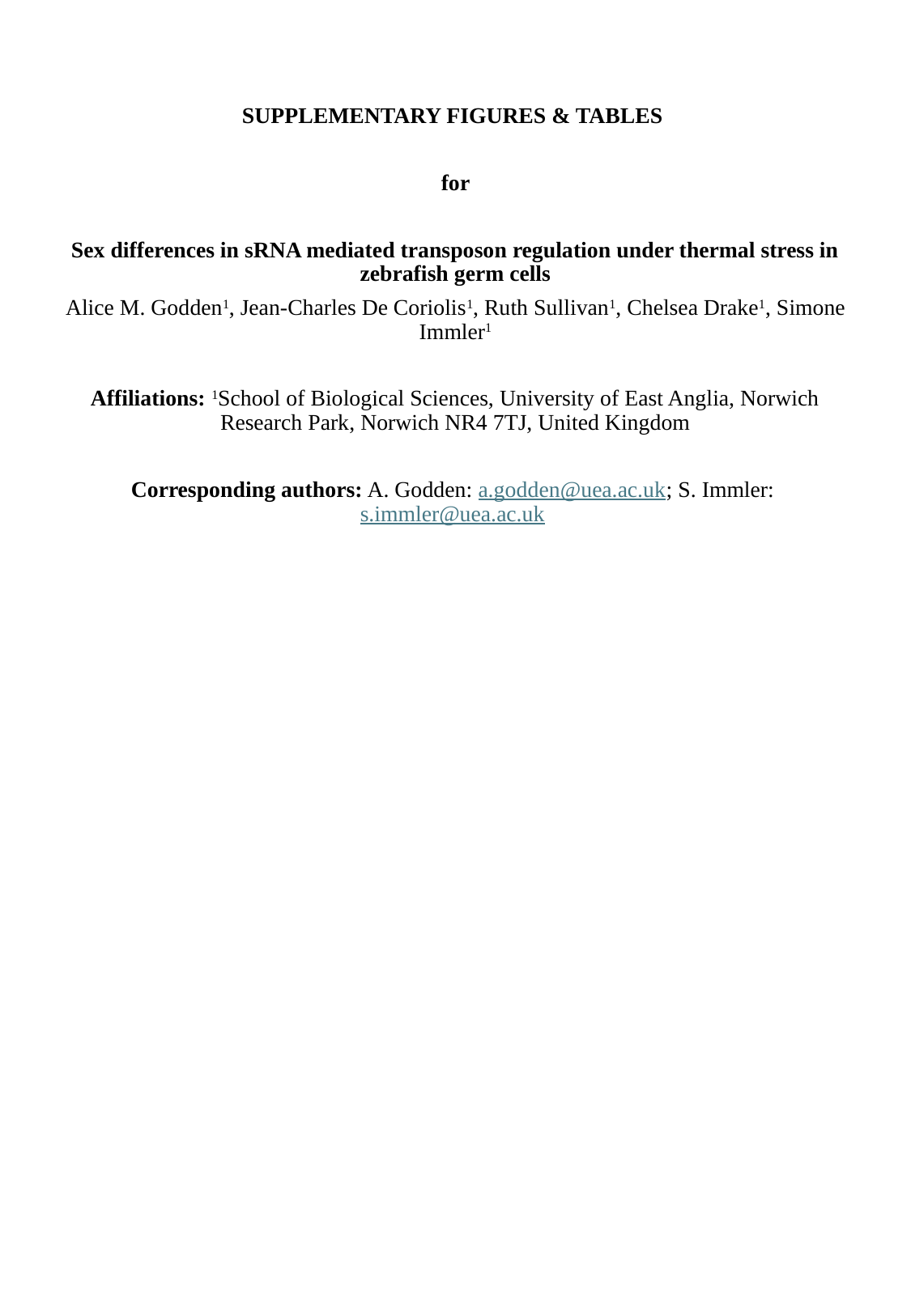

SUPPLEMENTARY FIGURES & TABLES
for
Sex differences in sRNA mediated transposon regulation under thermal stress in zebrafish germ cells
Alice M. Godden^1^, Jean-Charles De Coriolis^1^, Ruth Sullivan^1^, Chelsea Drake^1^, Simone Immler^1^
Affiliations: ^1^School of Biological Sciences, University of East Anglia, Norwich Research Park, Norwich NR4 7TJ, United Kingdom

### Slide 2
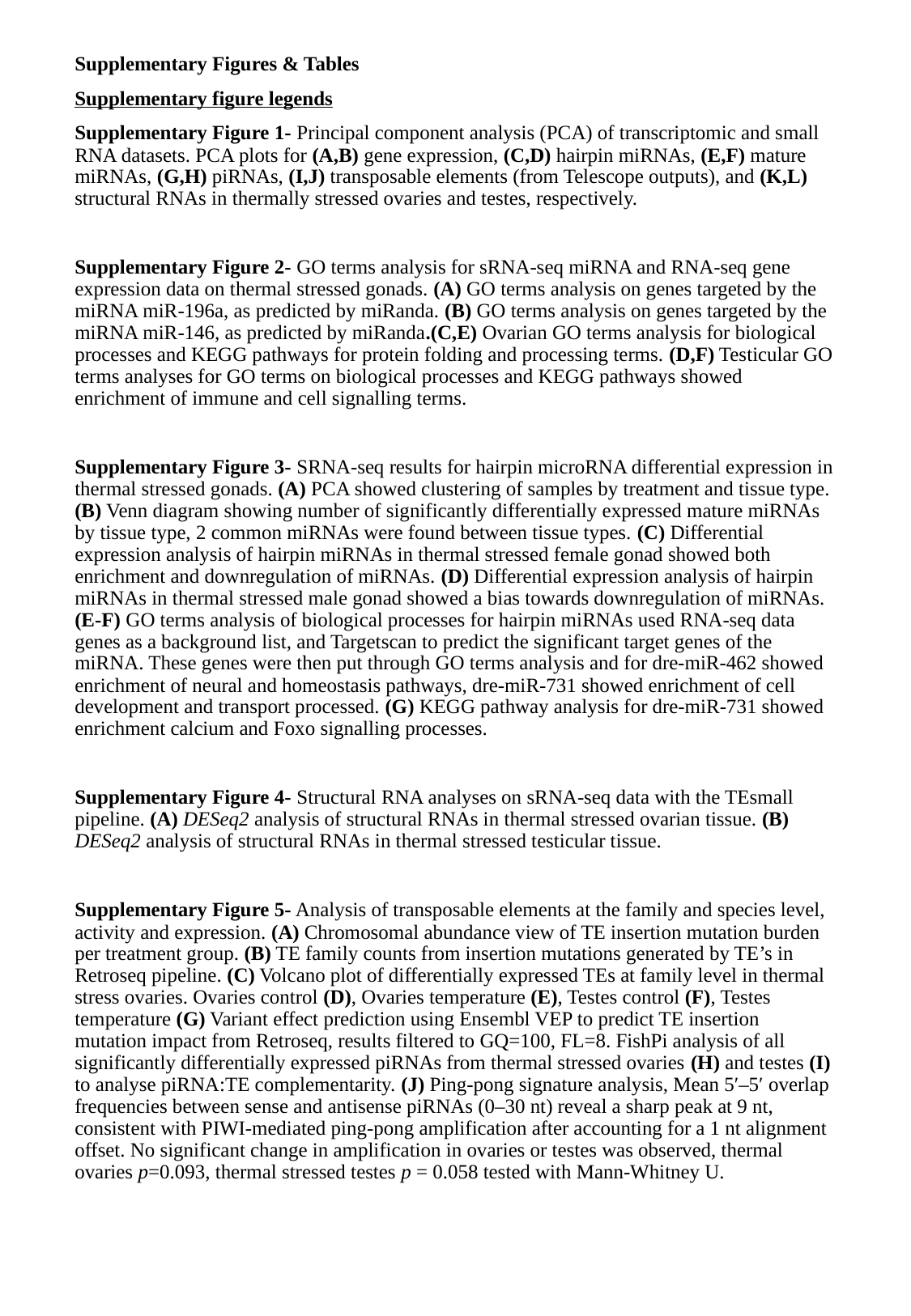

Supplementary Figures & Tables
Supplementary figure legends
Supplementary Figure 1- Principal component analysis (PCA) of transcriptomic and small RNA datasets. PCA plots for (A,B) gene expression, (C,D) hairpin miRNAs, (E,F) mature miRNAs, (G,H) piRNAs, (I,J) transposable elements (from Telescope outputs), and (K,L) structural RNAs in thermally stressed ovaries and testes, respectively.
Supplementary Figure 2- GO terms analysis for sRNA-seq miRNA and RNA-seq gene expression data on thermal stressed gonads. (A) GO terms analysis on genes targeted by the miRNA miR-196a, as predicted by miRanda. (B) GO terms analysis on genes targeted by the miRNA miR-146, as predicted by miRanda.(C,E) Ovarian GO terms analysis for biological processes and KEGG pathways for protein folding and processing terms. (D,F) Testicular GO terms analyses for GO terms on biological processes and KEGG pathways showed enrichment of immune and cell signalling terms.
Supplementary Figure 3- SRNA-seq results for hairpin microRNA differential expression in thermal stressed gonads. (A) PCA showed clustering of samples by treatment and tissue type. (B) Venn diagram showing number of significantly differentially expressed mature miRNAs by tissue type, 2 common miRNAs were found between tissue types. (C) Differential expression analysis of hairpin miRNAs in thermal stressed female gonad showed both enrichment and downregulation of miRNAs. (D) Differential expression analysis of hairpin miRNAs in thermal stressed male gonad showed a bias towards downregulation of miRNAs. (E-F) GO terms analysis of biological processes for hairpin miRNAs used RNA-seq data genes as a background list, and Targetscan to predict the significant target genes of the miRNA. These genes were then put through GO terms analysis and for dre-miR-462 showed enrichment of neural and homeostasis pathways, dre-miR-731 showed enrichment of cell development and transport processed. (G) KEGG pathway analysis for dre-miR-731 showed enrichment calcium and Foxo signalling processes.
Supplementary Figure 4- Structural RNA analyses on sRNA-seq data with the TEsmall pipeline. (A) DESeq2 analysis of structural RNAs in thermal stressed ovarian tissue. (B) DESeq2 analysis of structural RNAs in thermal stressed testicular tissue.
Supplementary Figure 5- Analysis of transposable elements at the family and species level, activity and expression. (A) Chromosomal abundance view of TE insertion mutation burden per treatment group. (B) TE family counts from insertion mutations generated by TE’s in Retroseq pipeline. (C) Volcano plot of differentially expressed TEs at family level in thermal stress ovaries. Ovaries control (D), Ovaries temperature (E), Testes control (F), Testes temperature (G) Variant effect prediction using Ensembl VEP to predict TE insertion mutation impact from Retroseq, results filtered to GQ=100, FL=8. FishPi analysis of all significantly differentially expressed piRNAs from thermal stressed ovaries (H) and testes (I) to analyse piRNA:TE complementarity. (J) Ping-pong signature analysis, Mean 5′–5′ overlap frequencies between sense and antisense piRNAs (0–30 nt) reveal a sharp peak at 9 nt, consistent with PIWI‑mediated ping‑pong amplification after accounting for a 1 nt alignment offset. No significant change in amplification in ovaries or testes was observed, thermal ovaries p=0.093, thermal stressed testes p = 0.058 tested with Mann-Whitney U.

### Slide 3
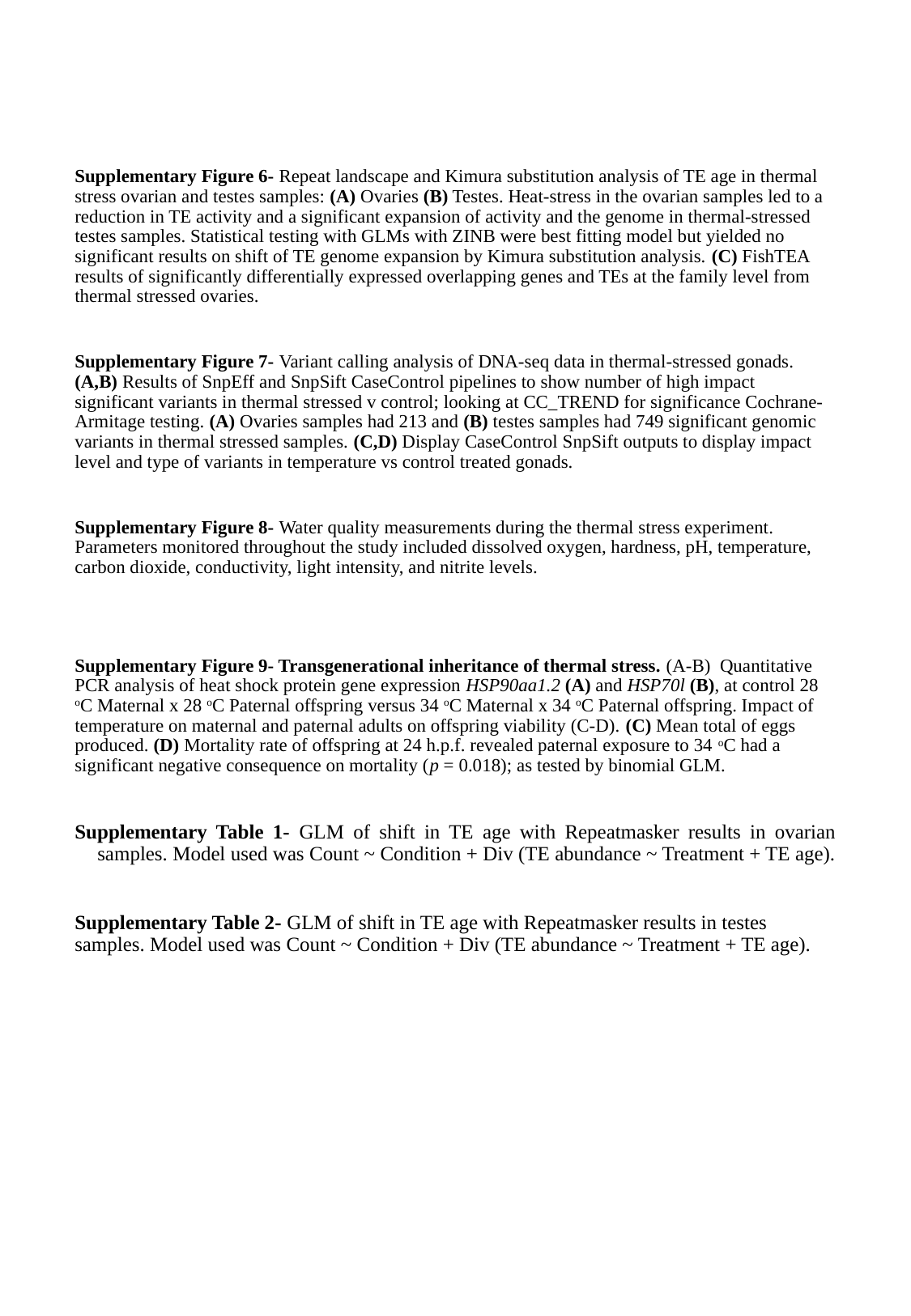

Supplementary Figure 6- Repeat landscape and Kimura substitution analysis of TE age in thermal stress ovarian and testes samples: (A) Ovaries (B) Testes. Heat-stress in the ovarian samples led to a reduction in TE activity and a significant expansion of activity and the genome in thermal-stressed testes samples. Statistical testing with GLMs with ZINB were best fitting model but yielded no significant results on shift of TE genome expansion by Kimura substitution analysis. (C) FishTEA results of significantly differentially expressed overlapping genes and TEs at the family level from thermal stressed ovaries.
Supplementary Figure 7- Variant calling analysis of DNA-seq data in thermal-stressed gonads. (A,B) Results of SnpEff and SnpSift CaseControl pipelines to show number of high impact significant variants in thermal stressed v control; looking at CC_TREND for significance Cochrane-Armitage testing. (A) Ovaries samples had 213 and (B) testes samples had 749 significant genomic variants in thermal stressed samples. (C,D) Display CaseControl SnpSift outputs to display impact level and type of variants in temperature vs control treated gonads.
Supplementary Figure 8- Water quality measurements during the thermal stress experiment. Parameters monitored throughout the study included dissolved oxygen, hardness, pH, temperature, carbon dioxide, conductivity, light intensity, and nitrite levels.
Supplementary Figure 9- Transgenerational inheritance of thermal stress. (A-B) Quantitative PCR analysis of heat shock protein gene expression HSP90aa1.2 (A) and HSP70l (B), at control 28 ^o^C Maternal x 28 ^o^C Paternal offspring versus 34 ^o^C Maternal x 34 ^o^C Paternal offspring. Impact of temperature on maternal and paternal adults on offspring viability (C-D). (C) Mean total of eggs produced. (D) Mortality rate of offspring at 24 h.p.f. revealed paternal exposure to 34 ^o^C had a significant negative consequence on mortality (p = 0.018); as tested by binomial GLM.
Supplementary Table 1- GLM of shift in TE age with Repeatmasker results in ovarian samples. Model used was Count ~ Condition + Div (TE abundance ~ Treatment + TE age).
Supplementary Table 2- GLM of shift in TE age with Repeatmasker results in testes samples. Model used was Count ~ Condition + Div (TE abundance ~ Treatment + TE age).

### Slide 4
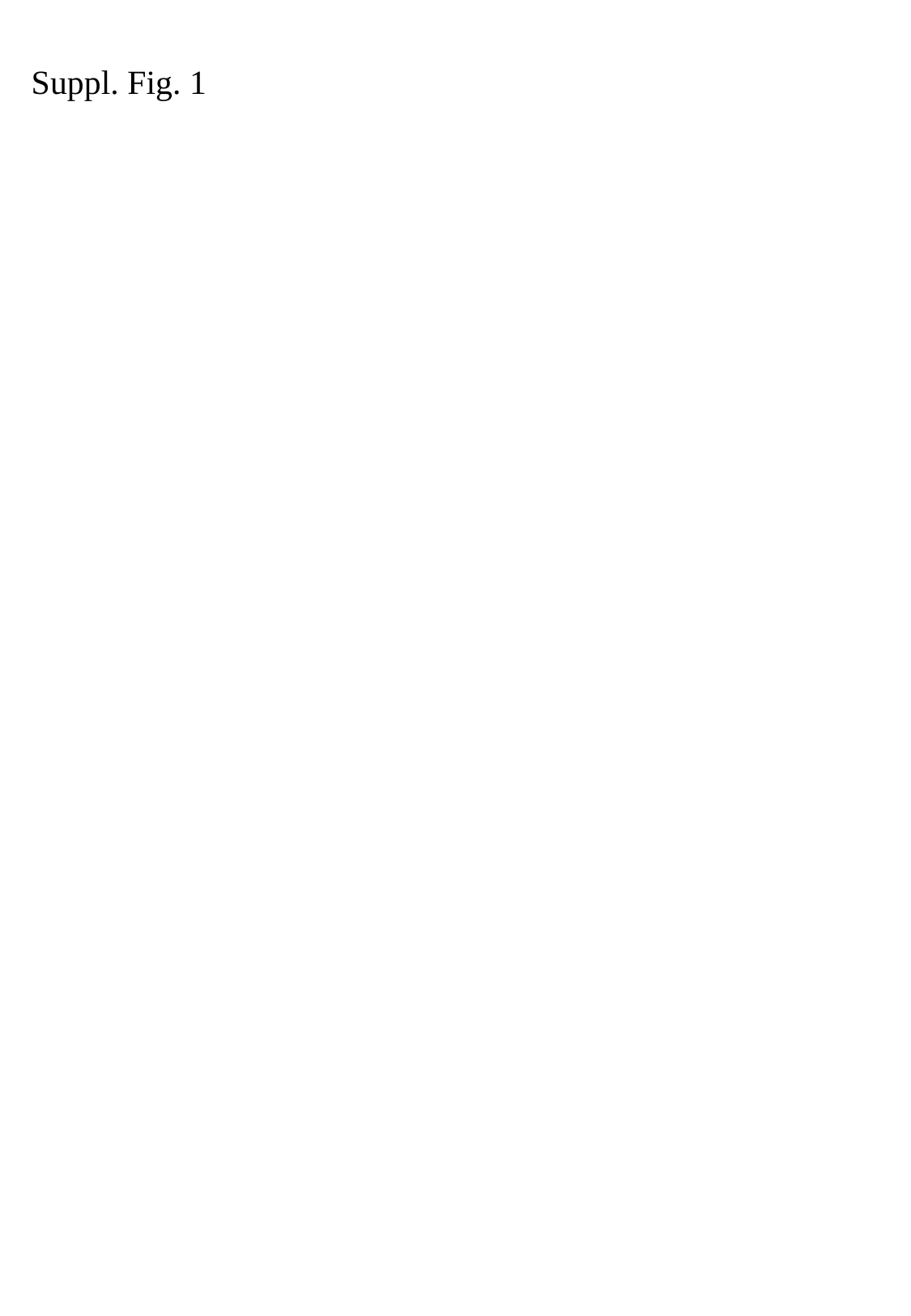

Suppl. Fig. 1

### Slide 5
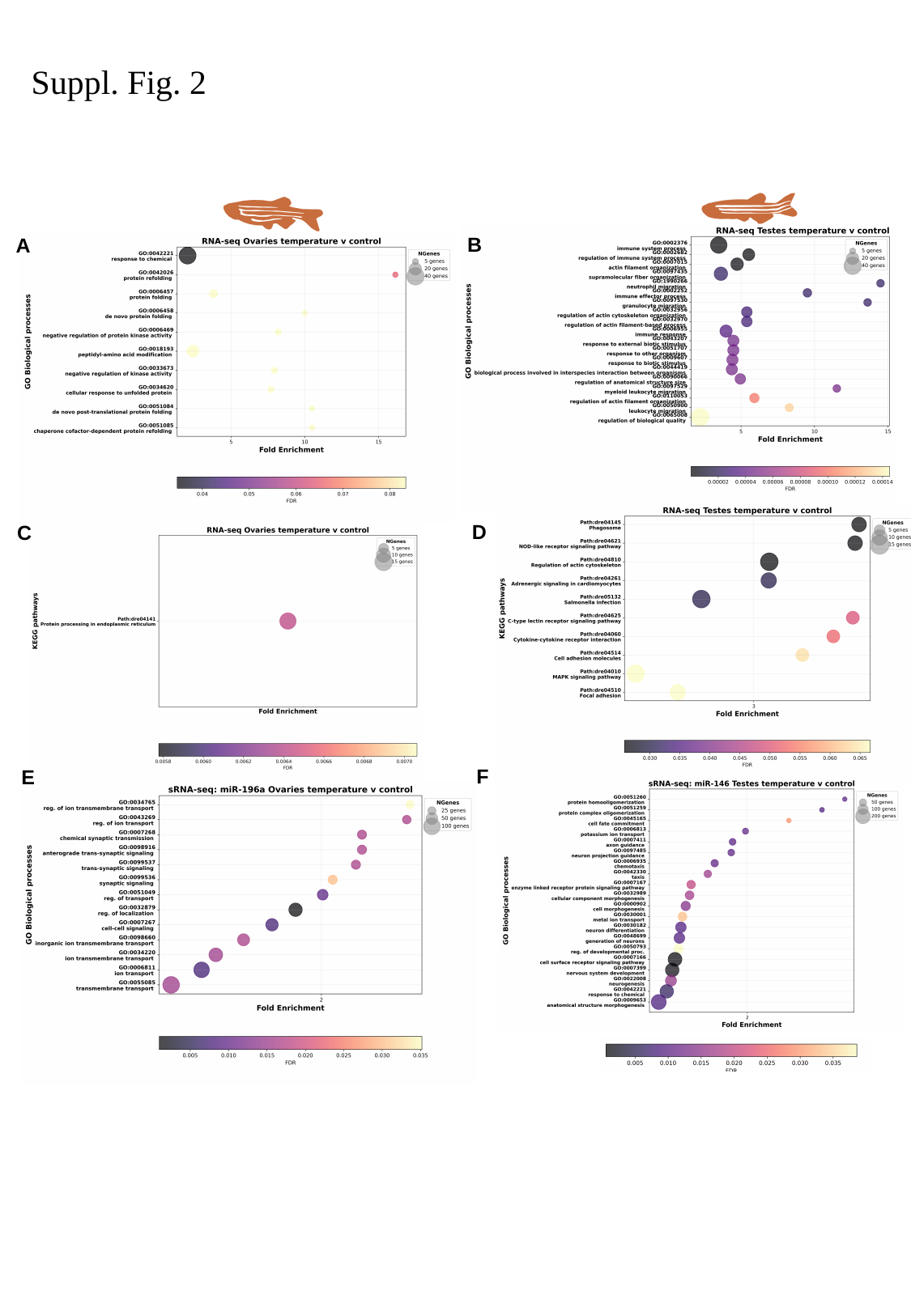

Suppl. Fig. 2
B
A
D
C
F
E

### Slide 6
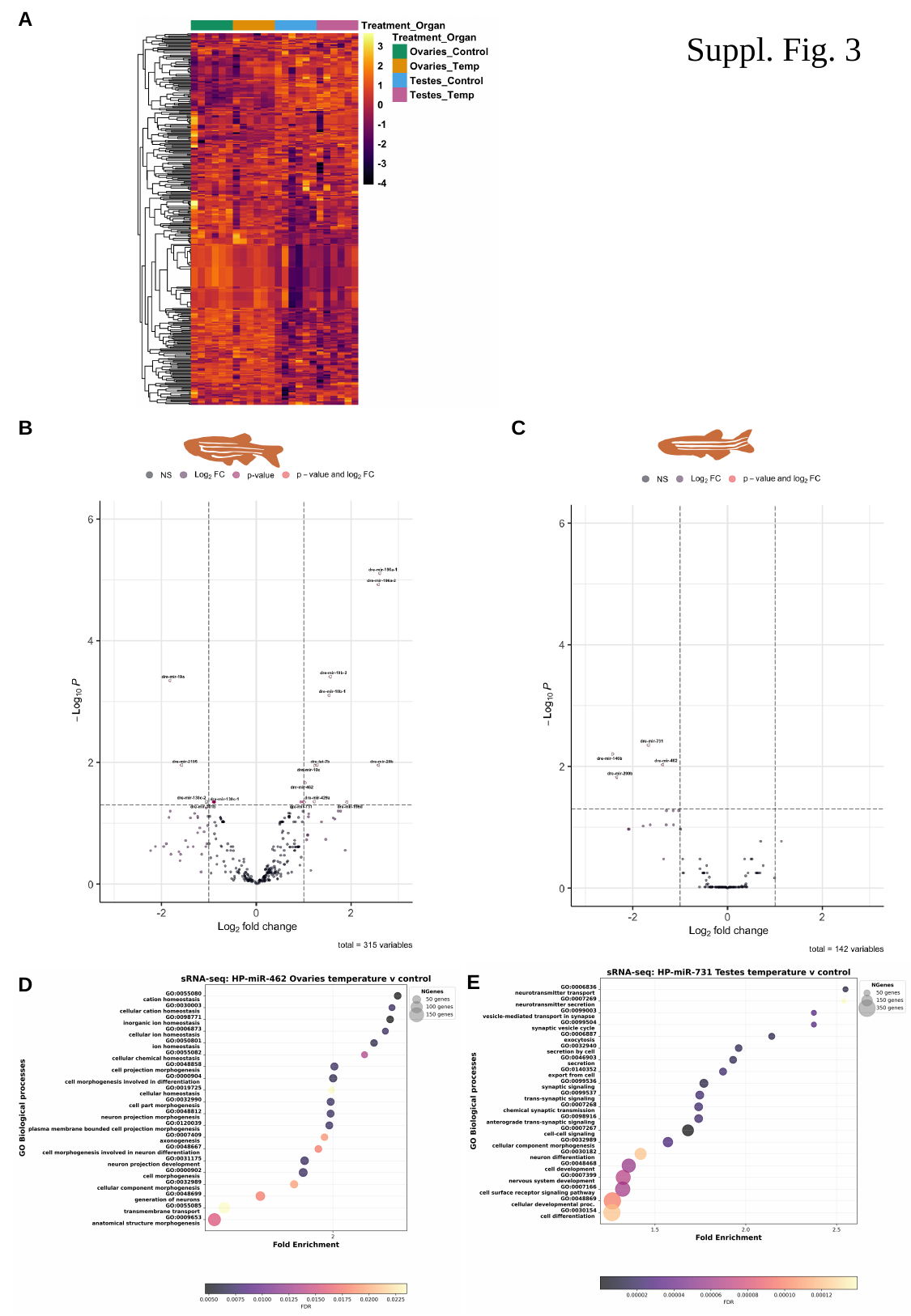

A
C
B
E
D
F
G
Suppl. Fig. 3

### Slide 7
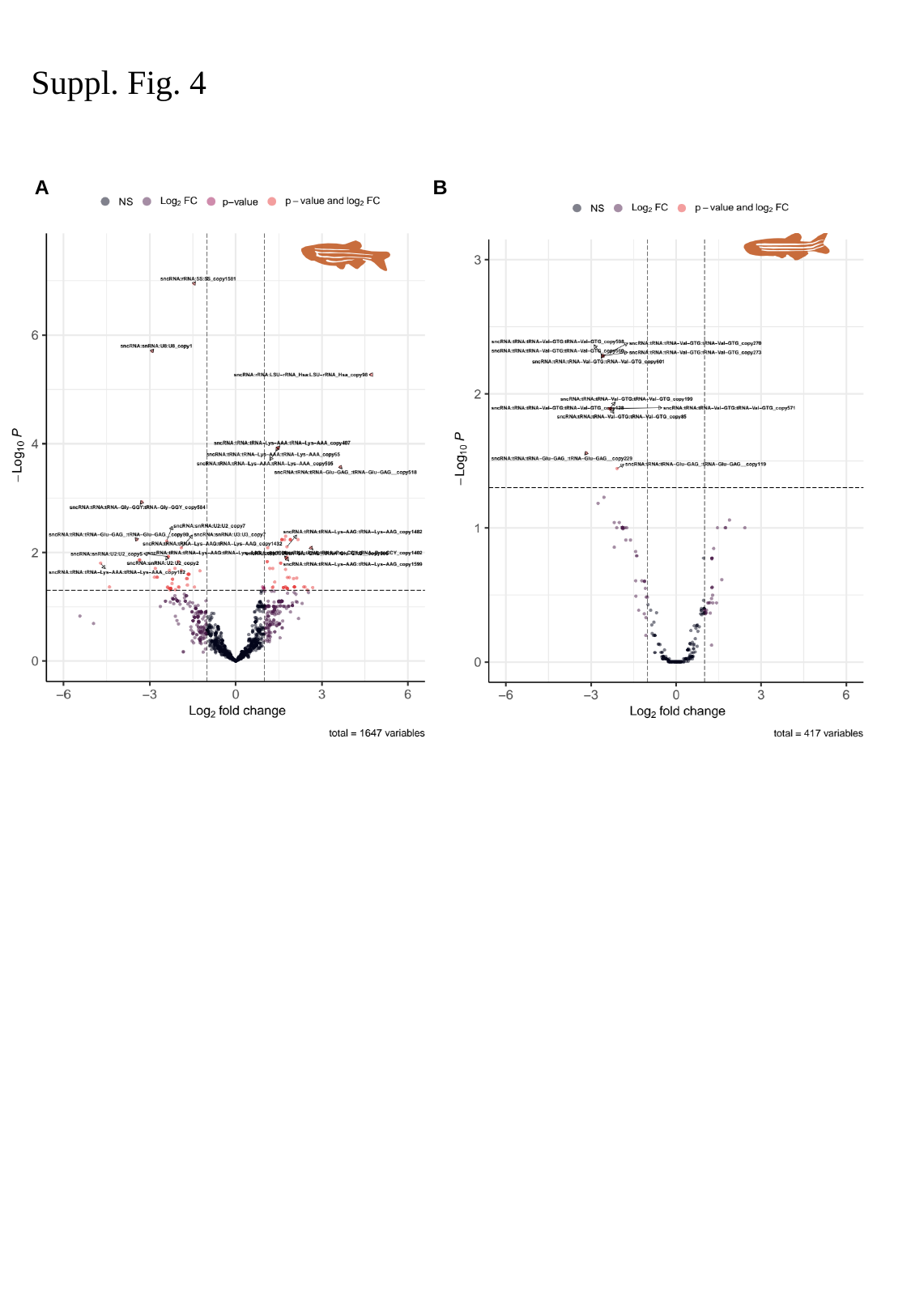

Suppl. Fig. 4
A
B

### Slide 8
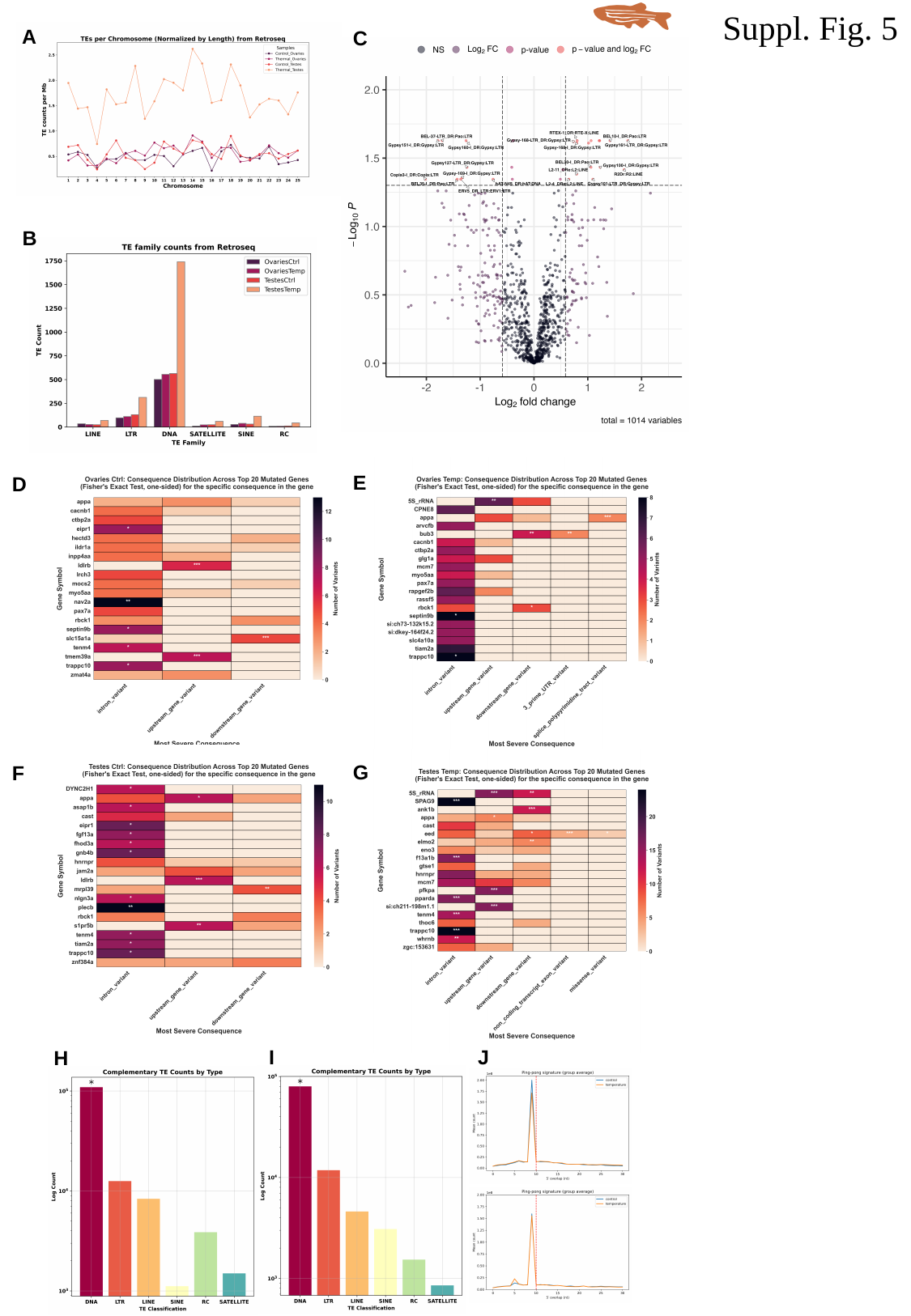

Suppl. Fig. 5
A
C
B
E
D
G
F
I
J
H

### Slide 9
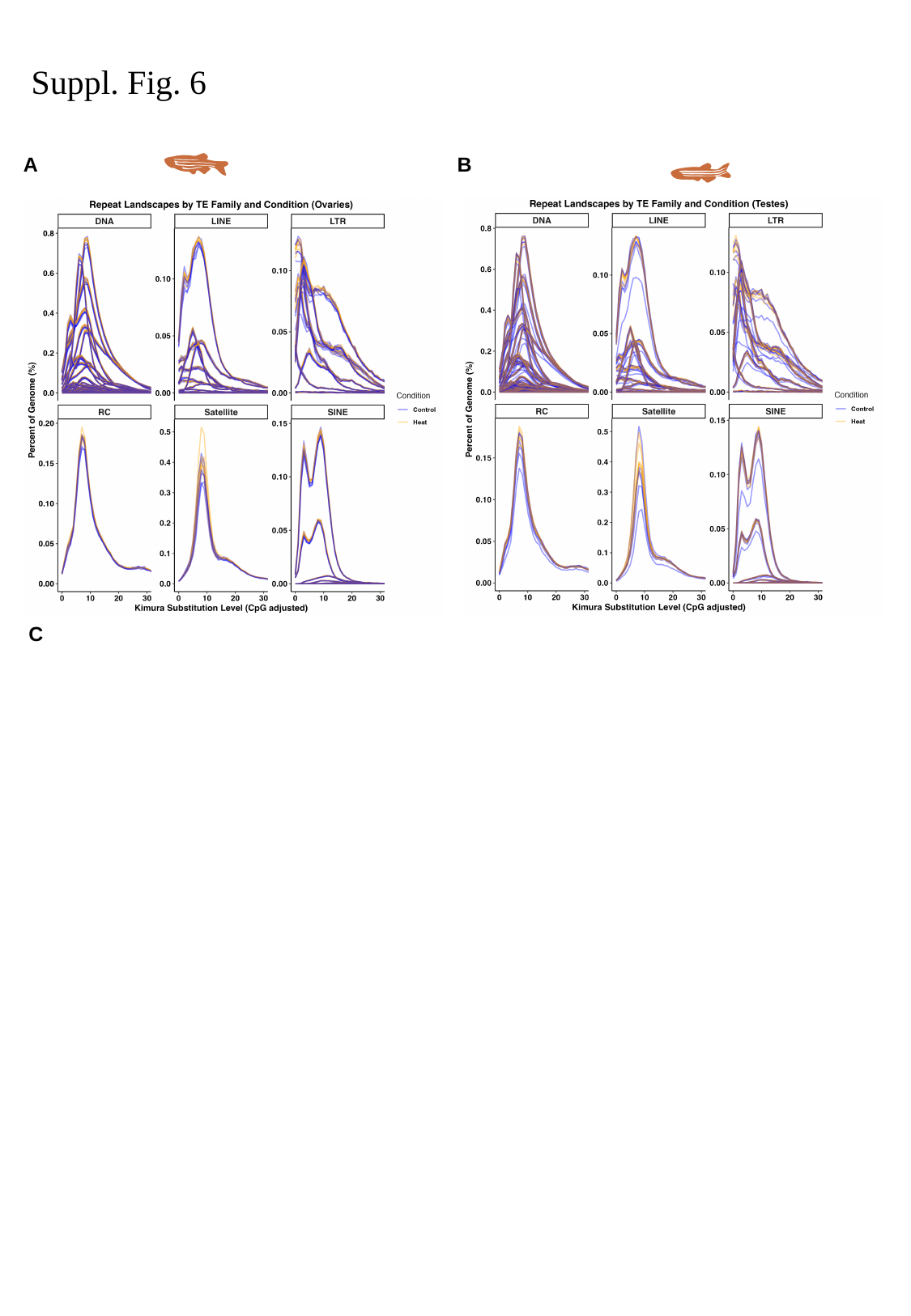

Suppl. Fig. 6
A
B
C

### Slide 10
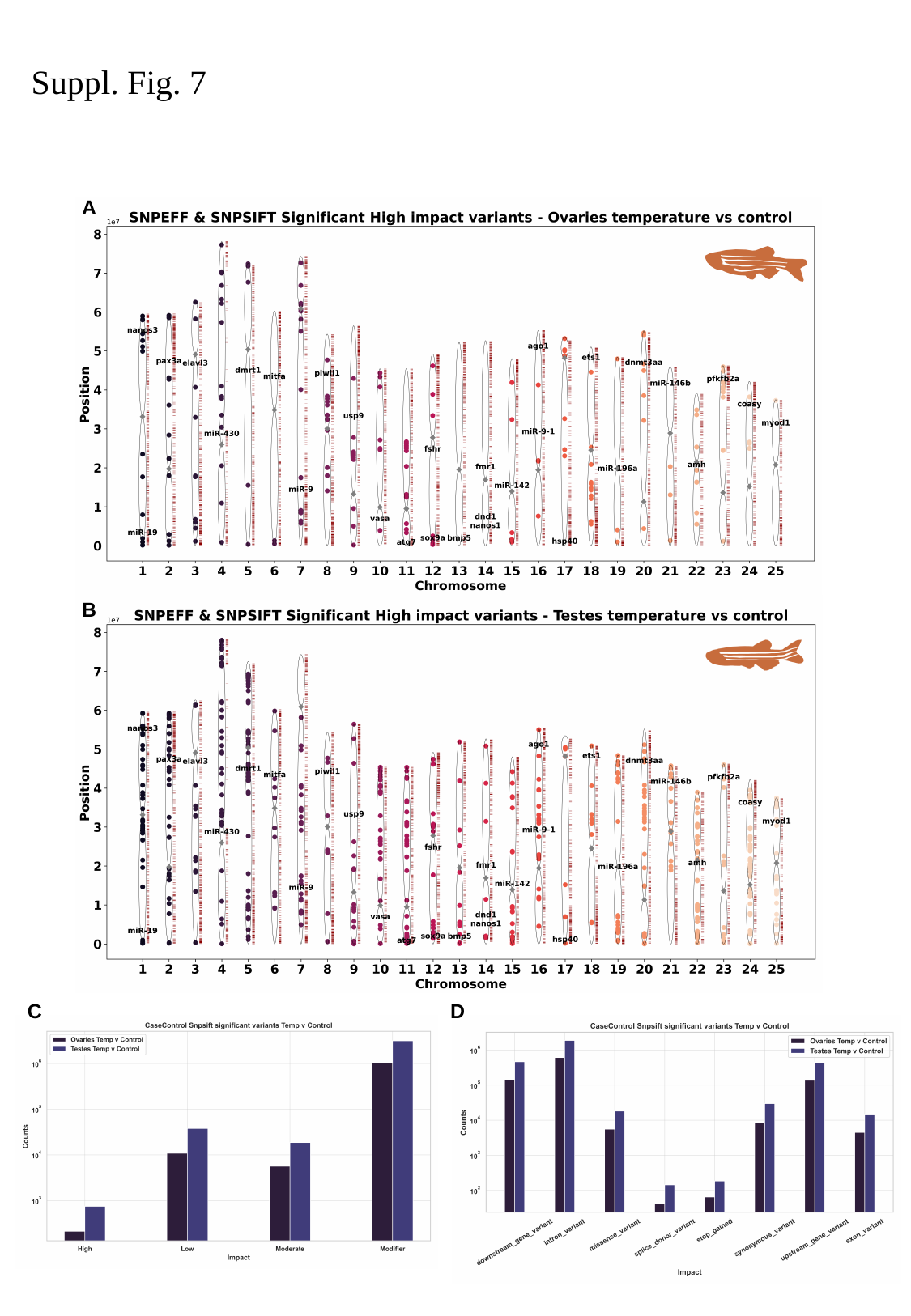

Suppl. Fig. 7
A
B
D
C

### Slide 11
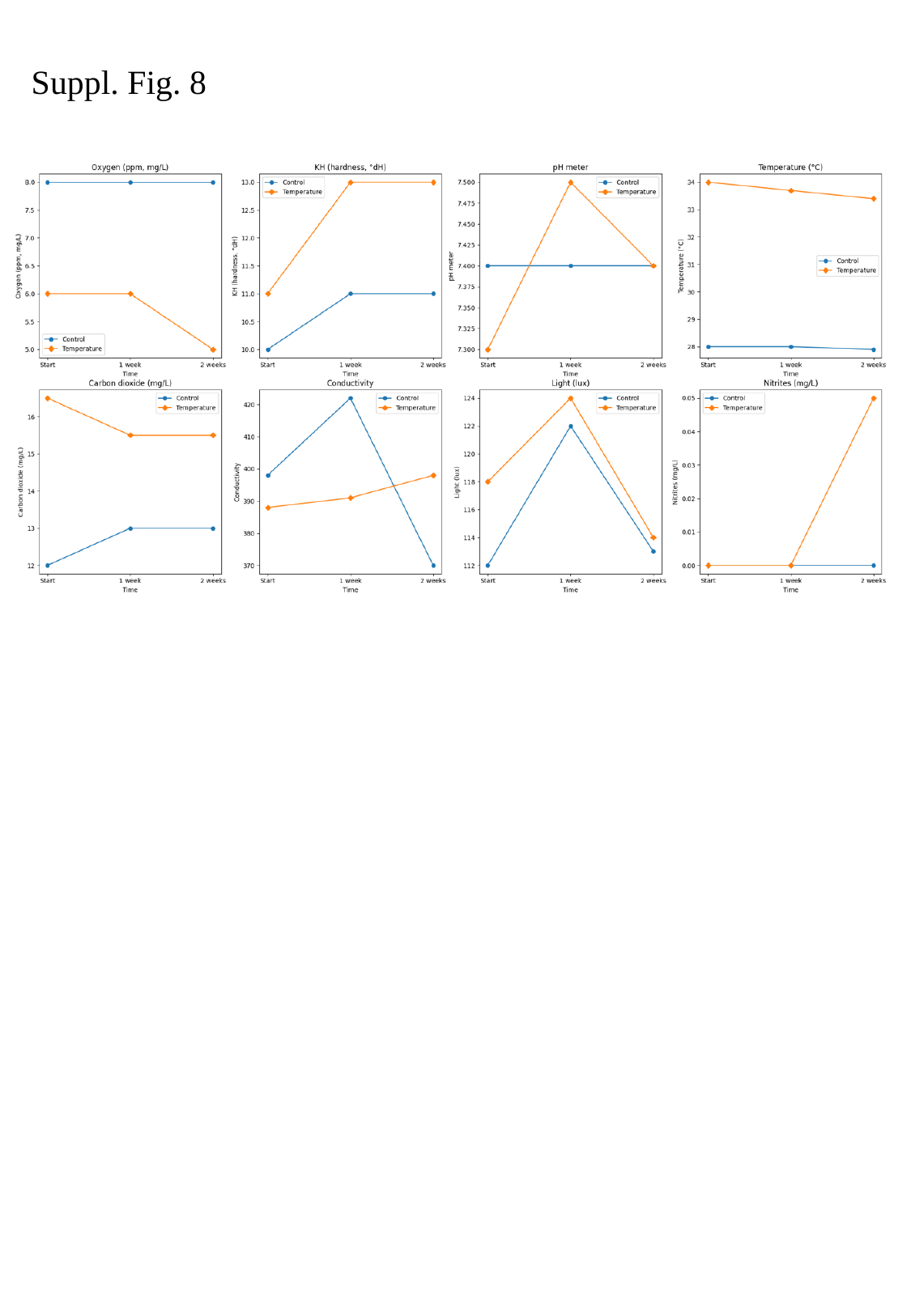

Suppl. Fig. 8

### Slide 12
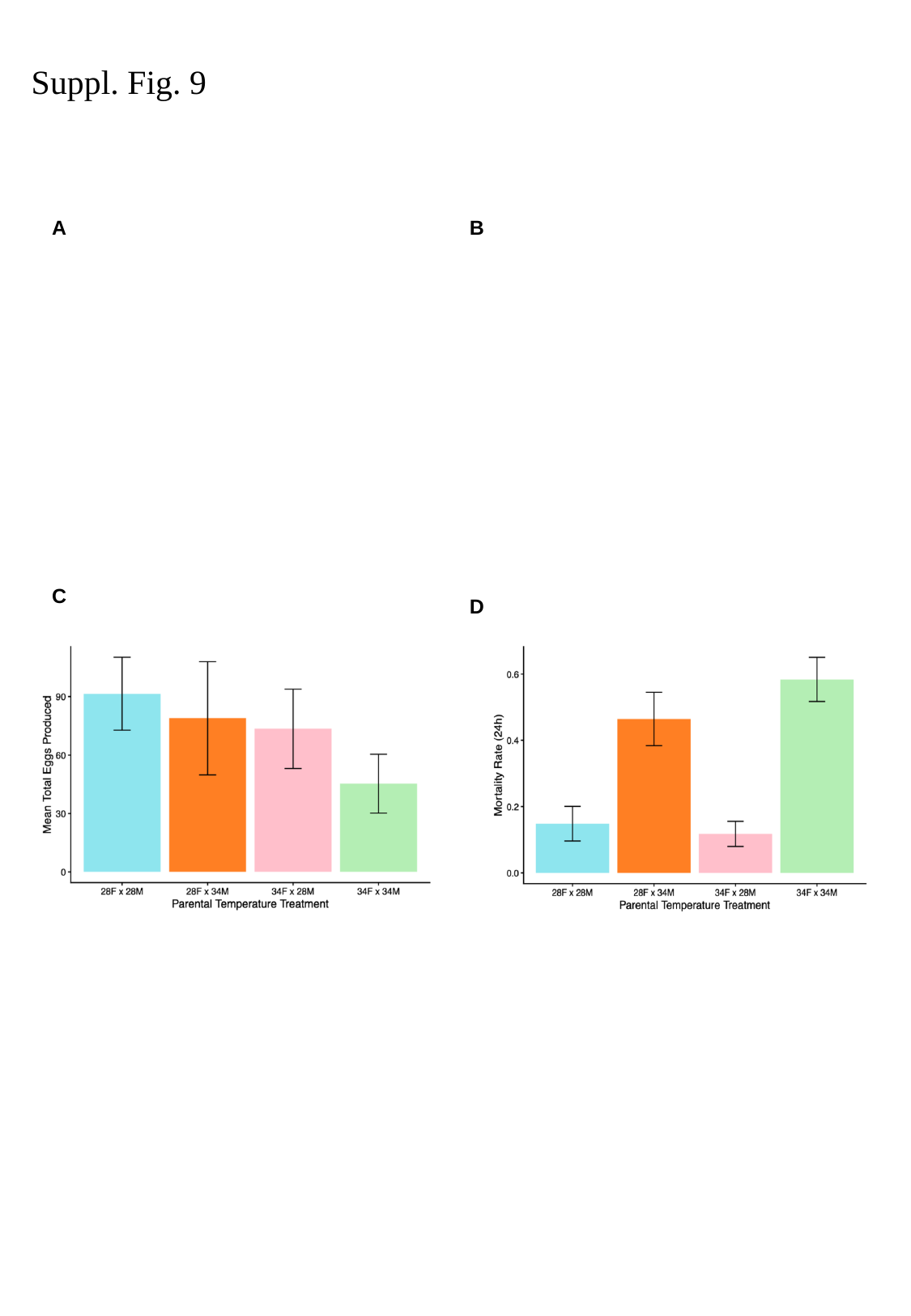

Suppl. Fig. 9
A
B
C
D

### Slide 13
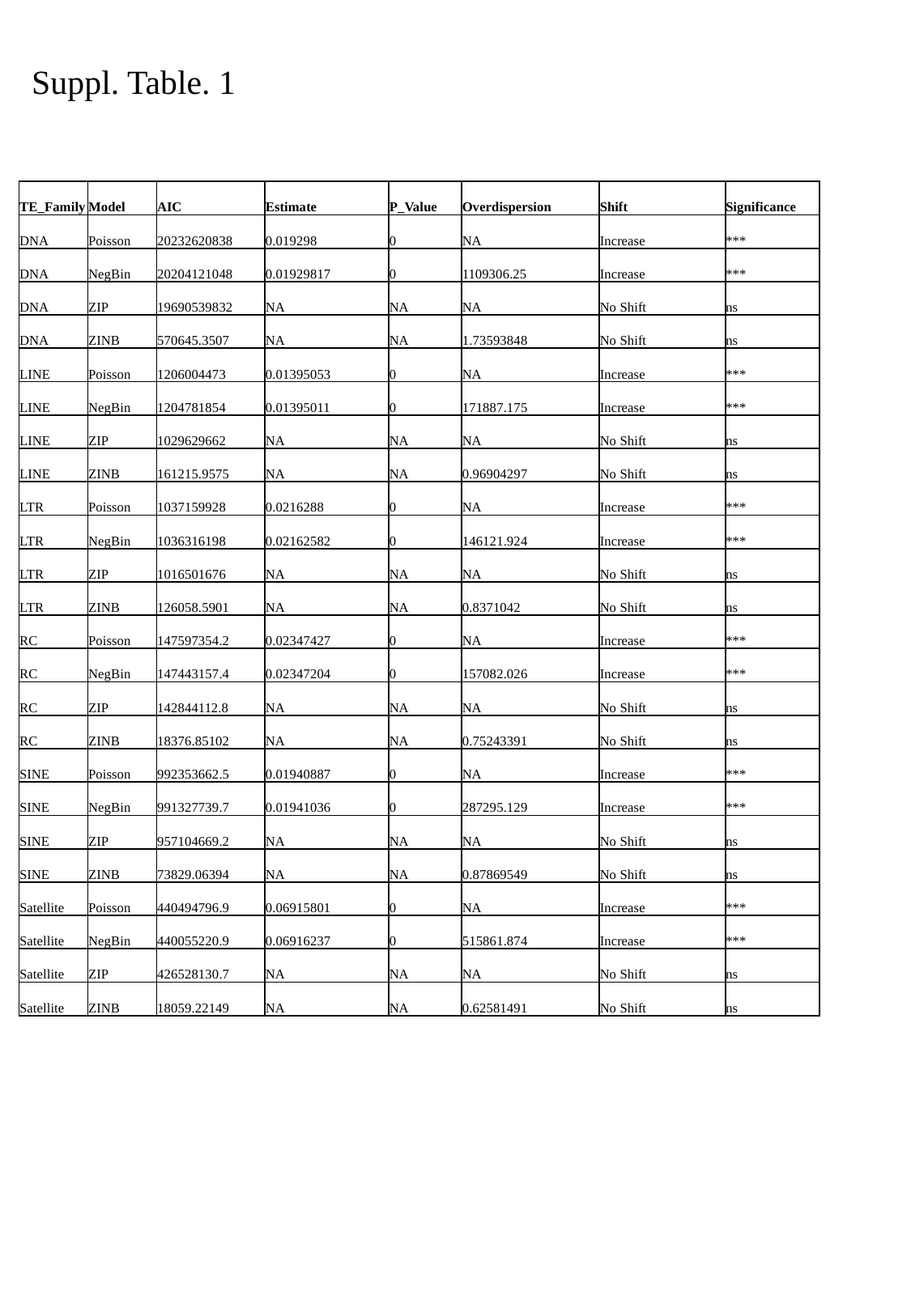

Suppl. Table. 1
| TE\_Family | Model | AIC | Estimate | P\_Value | Overdispersion | Shift | Significance |
| --- | --- | --- | --- | --- | --- | --- | --- |
| DNA | Poisson | 20232620838 | 0.019298 | 0 | NA | Increase | \*\*\* |
| DNA | NegBin | 20204121048 | 0.01929817 | 0 | 1109306.25 | Increase | \*\*\* |
| DNA | ZIP | 19690539832 | NA | NA | NA | No Shift | ns |
| DNA | ZINB | 570645.3507 | NA | NA | 1.73593848 | No Shift | ns |
| LINE | Poisson | 1206004473 | 0.01395053 | 0 | NA | Increase | \*\*\* |
| LINE | NegBin | 1204781854 | 0.01395011 | 0 | 171887.175 | Increase | \*\*\* |
| LINE | ZIP | 1029629662 | NA | NA | NA | No Shift | ns |
| LINE | ZINB | 161215.9575 | NA | NA | 0.96904297 | No Shift | ns |
| LTR | Poisson | 1037159928 | 0.0216288 | 0 | NA | Increase | \*\*\* |
| LTR | NegBin | 1036316198 | 0.02162582 | 0 | 146121.924 | Increase | \*\*\* |
| LTR | ZIP | 1016501676 | NA | NA | NA | No Shift | ns |
| LTR | ZINB | 126058.5901 | NA | NA | 0.8371042 | No Shift | ns |
| RC | Poisson | 147597354.2 | 0.02347427 | 0 | NA | Increase | \*\*\* |
| RC | NegBin | 147443157.4 | 0.02347204 | 0 | 157082.026 | Increase | \*\*\* |
| RC | ZIP | 142844112.8 | NA | NA | NA | No Shift | ns |
| RC | ZINB | 18376.85102 | NA | NA | 0.75243391 | No Shift | ns |
| SINE | Poisson | 992353662.5 | 0.01940887 | 0 | NA | Increase | \*\*\* |
| SINE | NegBin | 991327739.7 | 0.01941036 | 0 | 287295.129 | Increase | \*\*\* |
| SINE | ZIP | 957104669.2 | NA | NA | NA | No Shift | ns |
| SINE | ZINB | 73829.06394 | NA | NA | 0.87869549 | No Shift | ns |
| Satellite | Poisson | 440494796.9 | 0.06915801 | 0 | NA | Increase | \*\*\* |
| Satellite | NegBin | 440055220.9 | 0.06916237 | 0 | 515861.874 | Increase | \*\*\* |
| Satellite | ZIP | 426528130.7 | NA | NA | NA | No Shift | ns |
| Satellite | ZINB | 18059.22149 | NA | NA | 0.62581491 | No Shift | ns |

### Slide 14
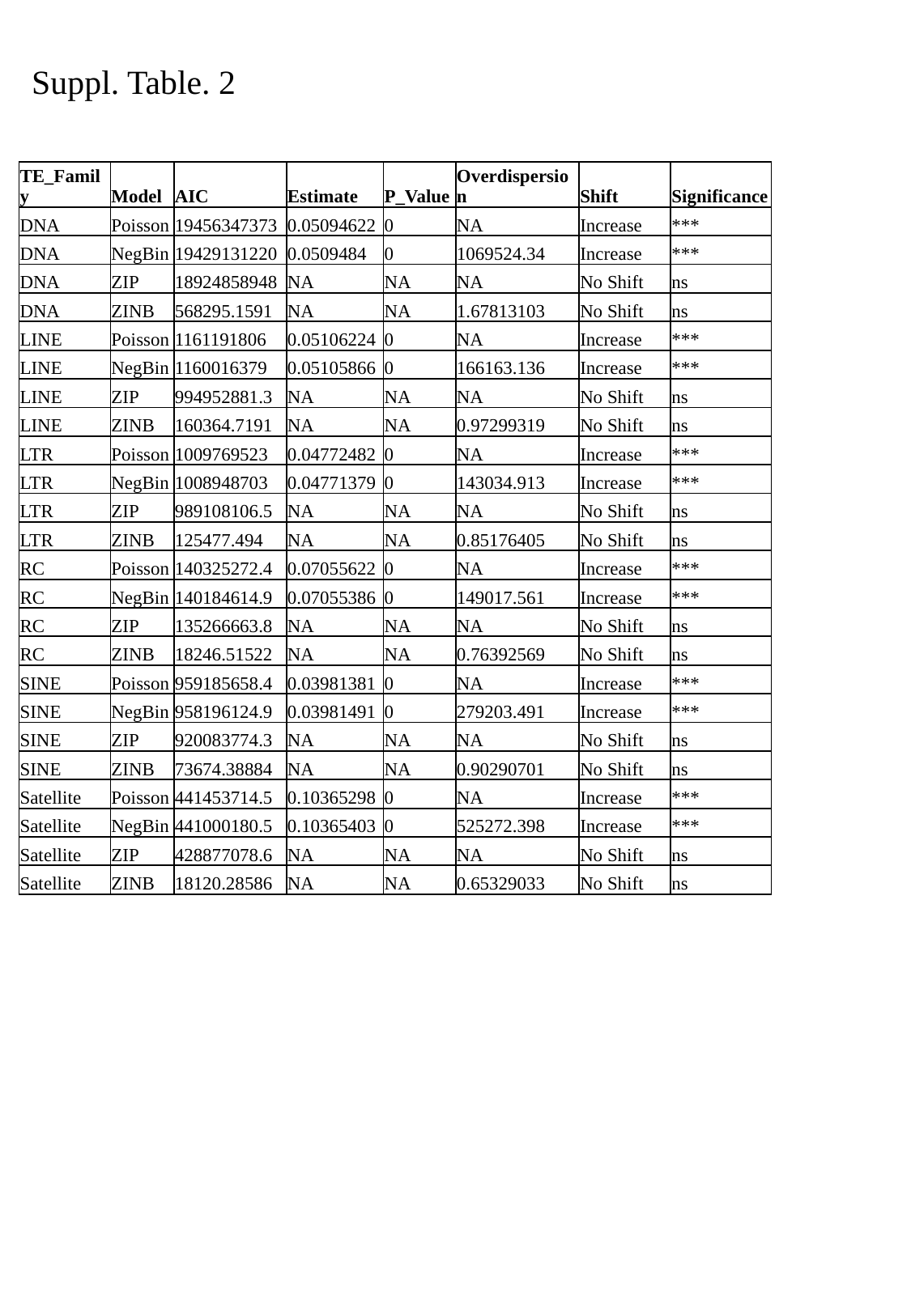

Suppl. Table. 2
| TE\_Family | Model | AIC | Estimate | P\_Value | Overdispersion | Shift | Significance |
| --- | --- | --- | --- | --- | --- | --- | --- |
| DNA | Poisson | 19456347373 | 0.05094622 | 0 | NA | Increase | \*\*\* |
| DNA | NegBin | 19429131220 | 0.0509484 | 0 | 1069524.34 | Increase | \*\*\* |
| DNA | ZIP | 18924858948 | NA | NA | NA | No Shift | ns |
| DNA | ZINB | 568295.1591 | NA | NA | 1.67813103 | No Shift | ns |
| LINE | Poisson | 1161191806 | 0.05106224 | 0 | NA | Increase | \*\*\* |
| LINE | NegBin | 1160016379 | 0.05105866 | 0 | 166163.136 | Increase | \*\*\* |
| LINE | ZIP | 994952881.3 | NA | NA | NA | No Shift | ns |
| LINE | ZINB | 160364.7191 | NA | NA | 0.97299319 | No Shift | ns |
| LTR | Poisson | 1009769523 | 0.04772482 | 0 | NA | Increase | \*\*\* |
| LTR | NegBin | 1008948703 | 0.04771379 | 0 | 143034.913 | Increase | \*\*\* |
| LTR | ZIP | 989108106.5 | NA | NA | NA | No Shift | ns |
| LTR | ZINB | 125477.494 | NA | NA | 0.85176405 | No Shift | ns |
| RC | Poisson | 140325272.4 | 0.07055622 | 0 | NA | Increase | \*\*\* |
| RC | NegBin | 140184614.9 | 0.07055386 | 0 | 149017.561 | Increase | \*\*\* |
| RC | ZIP | 135266663.8 | NA | NA | NA | No Shift | ns |
| RC | ZINB | 18246.51522 | NA | NA | 0.76392569 | No Shift | ns |
| SINE | Poisson | 959185658.4 | 0.03981381 | 0 | NA | Increase | \*\*\* |
| SINE | NegBin | 958196124.9 | 0.03981491 | 0 | 279203.491 | Increase | \*\*\* |
| SINE | ZIP | 920083774.3 | NA | NA | NA | No Shift | ns |
| SINE | ZINB | 73674.38884 | NA | NA | 0.90290701 | No Shift | ns |
| Satellite | Poisson | 441453714.5 | 0.10365298 | 0 | NA | Increase | \*\*\* |
| Satellite | NegBin | 441000180.5 | 0.10365403 | 0 | 525272.398 | Increase | \*\*\* |
| Satellite | ZIP | 428877078.6 | NA | NA | NA | No Shift | ns |
| Satellite | ZINB | 18120.28586 | NA | NA | 0.65329033 | No Shift | ns |
